# Diurnal variations in digestion and luminal flow determine microbial population dynamics along the human large intestine

**DOI:** 10.1101/2024.05.22.595260

**Authors:** Alinaghi Salari, Jonas Cremer

## Abstract

The human large intestine harbors a highly dynamic microbial ecosystem in which growing microbes regularly replenish biomass lost via feces. Understanding this population dynamics is biomedically important but remains a significant challenge due to rapid changes in microbial biomass and intestinal fluid flows. Leveraging experimental data on fluid turnover, nutrient supply, and microbial growth, we here derive a biophysical model of population dynamics. We show how the digestion of meals in batches triggers strong fluctuations in fluid movement and bacteria growth along the proximal large intestine. Comparing different model scenarios, we further analyze how the expandable nature of the proximal large intestine, the presence of a pouch-like cecum off major flow paths, and the periodic exit of luminal content via ‘mass movements’ are required in combination to maintain the high microbial population observed in the proximal large intestine. Since the microbial population undergoes several bottlenecks followed by rapid growth each day, the effective population size in the proximal large intestine is small, *N*_*e*_ ∼ 10^7^ − 10^11^, promoting the fast evolution of microbes. The diurnal fluctuations in flow also hamper the accumulation of slower-growing bacteria and lead to substantial variations in the uptake of fermentation products by the host. Our findings underscore the highly intertwined population dynamics of the gut microbiota, determined by the physics of fluid flow and growth with strong consequences for the microbiome and the host.

**Significance statement:** The density of microbial biomass within the large intestine is a key determinant of microbiome-host interactions and their impact on the human body. Studies with animal models have suggested highly intertwined population dynamics with strong variations of microbial densities over time and space. To elucidate the physiological drivers of these spatiotemporal dynamics along the human large intestine, we introduce a biophysical modeling framework that considers at its core the diurnal variation of digestion and fluid flow. Our analysis reveals how nutrient supply and the rapid movement of luminal content cause strong fluctuations in microbial biomass turnover throughout the day. The intertwined dynamics provide the host with ample mechanisms to control the microbiota, suppressing, for example, the emergence of slow-growing cross-feeding microbes.

## Introduction

The microbial population within the gut strongly influences human physiology and health, with its size and composition being key factors determining microbe-host interactions [1–5]. However, we poorly understand how microbial population dynamics emerge as a complex consequence of food intake, digestion, microbial growth characteristics, and the active movement of luminal content. This prevents more mechanistic insights into the functioning of this microbial ecosystem and its link to human physiology.

Along the small intestine, coordinated muscle contractions and high throughput of water lead to complex flow patterns that promote the efficient uptake of nutrients by the host while preventing a strong accumulation of microbes [6,7]. Gut microbes thus mostly reside in the large intestine, where they form a highly dynamic population. Microbial growth is primarily fueled by complex dietary carbohydrates escaping digestion along the upper digestive tract [8–11]. This growth compensates for the loss of microbial biomass regularly lost via feces. Importantly, this turnover is fast as around half of the microbial biomass is being replenished each day, with the exact turnover numbers depending on diet and the active movement of luminal contents (**SI Text 1**). Additionally, nutrient supply and luminal movements vary substantially throughout the day, tightly coupled to meal intake and digestion activity [12,13]. These diurnal variations and, more generally, the circadian rhythm of the host have been suggested to strongly impact the microbiota and its effect on host physiology [7,14–17]. In sum, microbial cell densities and the total population size of the microbiota in the large intestine thus emerge from a highly dynamic interplay between nutrient supply, microbial growth, and the active control and flow of luminal contents along the intestine with possibly strong consequences on the human host. How can we better understand this dynamic interplay and the different biological and physical factors shaping microbial population densities?

*In-vitro* and mathematical models are essential tools to better understand the microbial population dynamics along the human large intestine, given their potential to explore specific biophysical processes and the challenge of directly observing microbial densities within the human gut at relevant spatial and temporal resolutions. Different in-vitro models have been introduced to mimic aspects of microbial growth dynamics and turnover along the large intestine [18–21]. For instance, microfluidic approaches have been utilized to study the interaction of flow and turnover in spatial setups that emulate some characteristics of the intestine, including peristaltic movement, mixing, and interactions between microbiota and the epithelial layer [22–25]. However, in-vitro setups commonly focus on one or a few specific factors alone, and fixed experimental designs make it impractical to test the combined impact of multiple factors on microbial population dynamics. In parallel, mathematical modeling has thus been introduced to study the interplay of different physiological processes [19,23,26,27]. Mathematical models can also provide more mechanistic insights by accounting for the biological and physical fundamentals of involved processes, like the exponential nature of growth and the hydrodynamics of fluid transport. As such, they can be imperial in revealing the fundamental drivers of microbial population dynamics.

Most commonly, microbial growth is modeled in a series of well-mixed bioreactors or along a pipe-like structure with constant flow [20,28,29]. These modeling scenarios illustrate fundamental relationships between microbial growth, fluid turnover, and microbial density [30]. For example, they raise the question of how observed microbial densities can be maintained in the presence of high flow rates (**SI Text 2** and **Figs. S1 and S2)**. However, these models do not account for many key characteristics of the intestine that are experimentally well established, including strong diurnal fluctuations in flow and nutrient supply, motility features such as the gastro-colonic reflex, and geometrical properties like a cecum compartment. Here, we introduce an unprecedented modeling framework that integrates these key characteristics and also accounts for fundamental biophysical processes, including hydrodynamics, transport of luminal contents, microbial growth dynamics, and metabolite absorption kinetics. Our model thereby provides a range of mechanistic insights into microbial population dynamics along the large intestine and their strong fluctuations throughout the day.

## Results

### A Hydrodynamic Model of Microbial Growth along the Proximal Large Intestine

Supported by nutrient supply from the small intestine, microbial growth occurs mainly along the proximal large intestine, including the cecum and ascending colon. We thus focus our analysis on this intestinal segment. To investigate the growth dynamics of microbes, we set up a model that, following anatomy measurements (**Table S1**), emulates the proximal adult large intestine (LI) as a 30-cm long pipe-like geometry with a luminal volume of approximately 100 mL (illustration in **Fig. 1A, SI Text 3** and **Fig. S3**). We explicitly model the dynamics of nutrient concentrations, *c*_*n*_(*r, z, t*), and microbial densities, *c*_*b*_(*r, z, t*), in a 2-dimensional axisymmetric domain (coordinates *r* and z) over time *t* (**Fig. 1A**, Eqs. 1 and 2). Nutrients, primarily carbohydrates from diet not absorbed by the small intestine, enter with luminal fluids at the proximal side to support microbial growth (inflow, orange arrow in **Fig. 1A**). Microbes utilize these nutrients and grow with a growth rate depending on the local nutrient concentration and bound below a maximum rate, *λ* (Monod kinetics). Microbes and nutrients further move with the flow that emerges from the turnover of intestinal fluids and the mixing of fluids caused by the active contraction of the intestinal walls (i.e., gut motility) [31]. Supported by previous in-vitro studies we approximate this movement as an advection-diffusion process [26,32]. The drift velocity, *u* = (*u*_*r*_, *u*_*z*_), emerges from the net movement of intestinal fluids, which we account for by the solution of the

**Figure 1.**
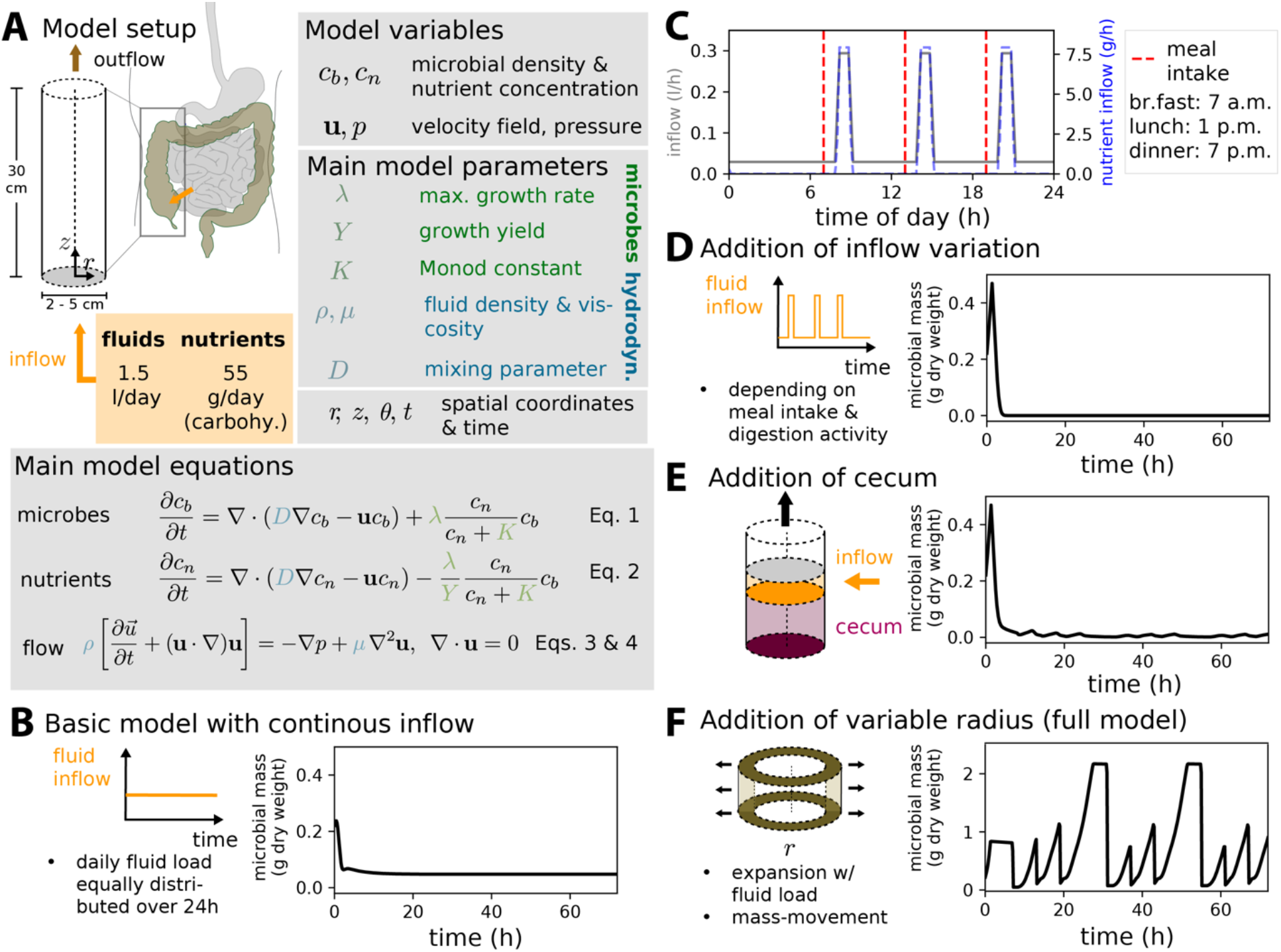
Modeling growth dynamics within the proximal end of the LI indicates dynamical volume changes required for promoting a stable microbial population. **(A)** Overview of the model setup, equations, and parameters. **(B)** Modeled daily variation of fluid load entering the cecum. **(C)** Basic model considering a pipe-like geometry and a constant daily fluid load entering (net inflow velocity of *u*_*z*_ *≈* 59 *μm/s* and nutrient concentration of 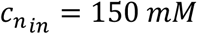). The graph shows the change in microbial biomass over 72 h with an initial dry microbe weight of ∼ 0.2 g, homogeneously distributed throughout the proximal LI. We consider three major meal intakes per day. Based on experimental observations, major peaks of fluid inflow and nutrient concentration occur at the cecum within 1 h post-meal intake (see **Fig. S4** for further details). **(D-F)** Extension of the basic model considering a diurnal varying inflow **(D)**, varying inflow and the presence of a cecum **(E)**, and varying inflow, a cecum, as well as regular diurnal expansion-contraction cycles of the radius, recapitulating volume control and mass movements along the proximal LI **(F)**. In all scenarios, the same initial conditions as in **C** are used. Used model parameters are provided in **Table S1** with the mixing strength of *D* = 10^−8^ *m*^2^*/s*. Further simulation details and results of these scenarios are discussed in **SI Text 6** and **Figs. S5-S9**.

Navier Stokes equation (**Fig. 1A**, Eq. 3) with appropriate boundary conditions (**Fig. S3**), while diffusion emulates the local mixing of content (see **SI Text 4**). Following the emerging flow dynamics, unabsorbed nutrients and microbes leave the system at the distal side (outflow, brown arrow in **Fig. 1A**). We use COMSOL Multiphysics to numerically solve the coupled system of equations using its built-in finite element method. As microbial growth parameters (highlighted in green in **Fig. 1A**), we use values we measured for primary fermenters that account for the vast majority of microbial biomass in the intestine [23]. Fluid-related parameters (highlighted in blue in **Fig. 1A**) follow from experimental studies (**SI Text 4 and 5**). We specifically use nutrient loads quantified for a British reference diet with an inflow of 1.5 L intestinal fluids per day [33,34]. All parameters used in this study are listed in **Table S1**.

### Model Scenarios of Increasing Complexity

The proximal human LI harbors a stable microbial population (**SI Text 7**) distinct from the small intestine [35]. To study how such a stable population can emerge, we start with the introduced basic model, which we subsequently extend in complexity to account for additional intestinal characteristics.

1. *Basic model with constant inflow:* We first consider a daily load of 1.5 L luminal fluids entering with a constant rate throughout the day (**Fig. 1B**). Notably, the radial variation of flow velocities in the pipe-like geometry, with flow velocities decreasing towards the pipe boundary, is not sufficient to overcome the high throughput of luminal contents and promote a stable microbial population. This is shown by the rapid decrease in population size when starting with a substantial microbial population in the system (**Fig. 1B**).
2. *Adding variations in fluid and nutrient inflow:* The fluid load entering the LI varies greatly over time, depending on food intake and digestion activities by the host. Measurements of water turnover [33] have shown that the inflow peaks approximately one hour after a major meal intake, while the basal inflow rate in the absence of food intake and digestion is very low (data analyzed in **Fig. S4**), suggesting a highly regulated fluid flow passing the ileocecal valve. We here specifically consider a scenario with three major peaks in inflow corresponding to three major meal intakes following measurements [33] in diet-controlled studies (**Fig. 1C**). Importantly, this strong variation in flow still does not support observed higher microbial densities, even throughout the night when the inflow is at its low basal rate. The total population size falls to very low levels (**Fig. 1D**).
3. *Adding cecum:* Next, we consider whether a dead-end, pouch-like cecal compartment can help stabilize microbial populations. To account for the cecum of the human adult, we specifically relocate the ileocecal valve from which nutrients enter to the side of the simulation domain at a 2 cm distance from the proximal end (highlighted in orange in **Fig. 1E, SI Text 3** and **Eq. [S16]**). Importantly, while a higher microbial population persists in the cecum, the addition of a cecum alone is not sufficient to ensure the higher microbial density along the ascending colon, and an initially high abundance of microbes is still being washed out over time in contrast to observations (**Fig. 1E**).
4. *Adding dynamical expansion-contraction of the colonic volume:* Scintigraphy, magnetic resonance imaging measurements, and intervention studies that control fluids and contents reaching the LI indicate substantial variations in the colonic volume with intestinal content being moved in batches [36–38]. These processes are highly controlled, coordinated with digestion and sleeping/awakening periods [39], and commonly referred to as “mass movement”. The most prominent example is the gastrocolic reflex with motility and peristaltic motion depending on the stretch of the stomach wall [40], such that mass movements occur before digested food reaches the LI (**SI Text 5**). To account for such a change in the intestinal volume and the rapid move of colonic contents, we extend the model to allow for dynamical changes of the intestinal radius *r* within the observed range of ∼1 ≤ *r* ≤ ∼2.7 *cm* (**Fig. 1F**; the time-dependent colonic radius is shown **in Fig. 2A** below). We specifically model the radius to increase gradually with fluid inflow from the small intestine, while a mass movement leads to a rapid shrinkage in radius and thus to the rapid movement and outflow of luminal content. Accounting for the gastrocolic reflex [41], we specifically implement mass movements after every meal intake and before substantial fluid loads reach the LI (**See SI - Eqs. [S3]-[S8])**. Crucially, the dynamic adjustment of the radius strongly influences microbial growth and turnover dynamics. As a result, high microbial densities emerge and remain stable (**Fig. 1F**), consistent with observed densities (see **SI Text 7** for a detailed discussion of numbers) [42,43]. In sum, our model connects nutrient supply, flow dynamics, and microbial growth to simulate microbial population dynamics within the proximal LI. Given the high inflow of luminal contents, an expandable volume of the proximal LI is required to ensure the observed high microbial densities within that intestinal section.

**Figure 2.**
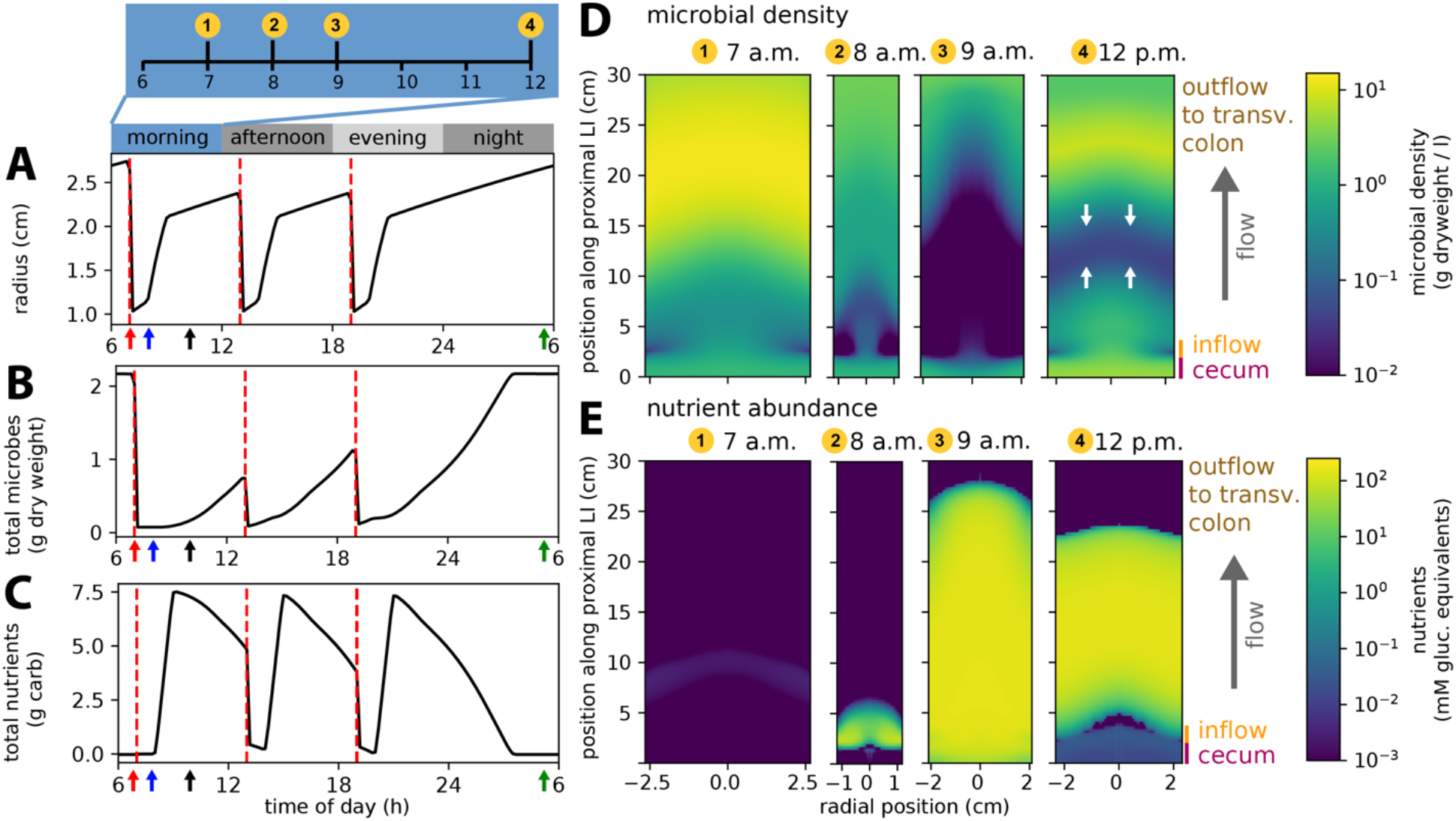
Spatiotemporal growth dynamics. **(A-C)** Temporal variation in radius and the abundance of microbes and nutrients within the proximal LI. The red, blue, black, and green arrows show the timing of the initiation of meal intake and subsequent mass movements, major fluid inflow to the cecum, significant microbial growth phases, and complete nutrient depletion, respectively. **(D, E)** Spatial variation of microbial densities and nutrient concentrations at selected time points. The white arrows in (D, E) indicate the two major directions of microbe repopulation in the ascending colon, influenced by a combined effect of growth and mixing. The presented results reflect a stable microbial population state with recurring daily variations. Full temporal dynamics over 72 h shown in **Fig. S6**. Model parameters provided in **Table S1**, with mixing strength *D* = 10^−8^ *m*^2^*/s*. Time-lapsed dynamics shown in **Video S1**.

### Diurnal Variations in Growth and Population Size

To better understand the emerging population dynamics and its diurnal variation with meal intake and digestion, we next characterize the full model with volume extension and mass movements in more detail. We specifically study the daily variation in radius, microbial densities, and nutrient contents over a 24-hour period, following the attainment of a high, albeit fluctuating, microbial population level after a sufficiently long (2-4 d) simulation time (**Fig. 2**). With meal intake at 7 a.m., mass movements lead to a rapid decrease in the radius (**Fig. 2A**, red arrow), pushing microbes out of the proximal LI and leading to a drop in the total number of microbes (**Fig. 2B**, red arrow). This prepares the colon for the subsequent intake of large amounts of fluids, which enter approximately one hour after meal intake, following digestion in the stomach and small intestine. As the radius begins to increase (**Fig. 2A**, blue arrow), more nutrients enter the intestine, supporting growth (**Fig. 2C**, blue arrow). During this phase, no major outflow occurs [44] (in the simulation, the outflow is assumed to be zero), microbes grow, and the population size increases significantly over time (**Fig. 2B**, black arrow). Similar dynamics, including mass movements and subsequent growth phases, repeatedly occur during the rest of the day following two additional meal intakes at 1 p.m. and 7 p.m.. After the third meal and during the night, when mass movements are less common (no mass movement in the simulations), the microbial population size can increase even higher with all available nutrients being consumed (**Fig. 2C**, green arrow).

To further illustrate these dynamics, we show the spatial variation of microbial density at four time points in the morning (**Fig. 2D**). Notably, the decrease in radius pushes luminal contents with high microbial densities out of the proximal intestine, leaving relatively low densities behind (time points 1 and 2). Local microbial densities become even more depleted with the strong inflow of fluids and nutrients (time point 3). Subsequently, growth leads to a strong increase in microbial densities (time point 4), with microbes from the cecum and distal part repopulating the entire ascending colon (white arrows). The concentration of nutrients shows a reverse trend. Starting with low concentrations, the influx of nutrients after meal intake leads to a strong increase in nutrient concentrations, followed by a subsequent decrease when the density of nutrient-consuming microbes increases (**Fig. 2E**). The full dynamics over a time span of 24 hours is shown in **Video S1**. Overall, these findings emphasize highly intertwined population dynamics with strong variations in growth and population size throughout the day, emerging from a tight interplay between meal intake, intestinal fluid turnover, and microbial growth.

### Variations in Microbial Growth Rates

To emphasize the strong consequences of this interplay, we next discuss how variations in the maximum growth rate, *λ*, affect population dynamics. The ability of microbial cells to grow largely depends on the coordination of major cellular processes to maintain metabolism and synthesize new biomass, resulting in strong variations of growth rates across species [45,46]. For example, primary fermenters utilize carbohydrates and commonly grow rapidly, with doubling times as short as 30 minutes (*λ*≳ 1.4 1*/h*) [9]. In contrast, cross-feeding species that use various metabolic strategies to grow anaerobically on fermentation products, including sulfate reduction and methanogenesis [47,48], commonly grow up to five times slower [49]. We, therefore, run simulations for a range of growth rates between *λ* = 0.2 1*/h* and 1.5 1*/h*. Crucially, the flow conditions along the intestine serve as an efficient “growth filter,” with only sufficiently fast-growing microbes being able to overcome the strong flow and establish a substantial population within the LI (**Figs. 3 and S10**). As such, flow hampers the accumulation of slow-growing cross-feeding species that could otherwise consume a high fraction of the fermentation products primary fermenters release. In summary, these results indicate that population dynamics vary substantially with encountered flow conditions, highlighting the host’s ability to shape microbial population dynamics and composition.

**Figure 3.**
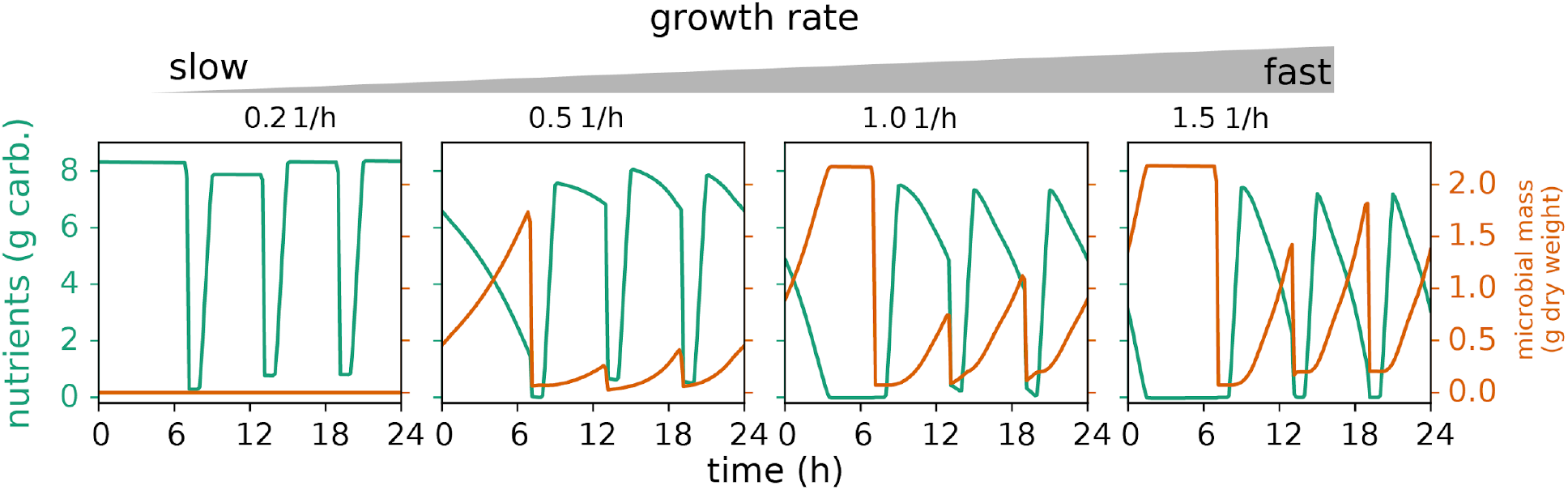
Dependence of microbial abundance on growth behavior. Temporal variation in microbial abundance within the proximal LI for different growth rates, *λ*, ranging from 0.2 to 1.5 1*/h* (corresponding to doubling times between ∼ 0.5 *h* and ∼ 3.5 *h*) with a mixing strength of *D* = 10^−8^ *m*^2^*/s*. Other simulation parameters are listed in **Table S1**. See **Fig. S10 for** corresponding kymographs showing spatiotemporal density variations.

### Cecum Promotes Growth and Determines Effective Population Size

The number of cells forming a microbial population strongly shapes the outcome of evolution. Previous work has highlighted the important role of flow dynamics in setting effective population sizes and, thus, evolutionary dynamics in gut environments [32,50]. We hence investigate in more detail the consequences of the discussed population dynamics on the number of microbes in the proximal LI. Notably, the microbial population in the cecum promotes growth along the proximal LI after meal intake, as illustrated by a kymograph showing microbial density (**Fig. 4A**) with a rapid drop of microbial density after a meal intake (gray arrow) followed by a recovery from the cecum (black arrow). Furthermore, no substantial microbial population remains in simulations without a cecum (**Fig. 4B** dashed line, and **Figs. S9**). The cecal microbial population is thus essential in shaping the population dynamics along the entire proximal LI. The population size in the cecum *N*_*cec*_ varies strongly over time (**Fig. 4B**, solid line), reaching the lowest values 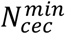 when the inflow from the small intestine peaks after meal intake and before the cecal population starts repopulating the proximal LI (**Fig. 4B**, markers). With evolution strongly determined by regular bottlenecks in population size [32,50,51], and the cecal population seeding growth in the ascending colon, 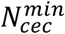 might be considered as the effective population size *N*_*e*_ of the system, varying between 10^9^ and 10^11^ cells, much lower than the *≈* 4 × 10^13^ microbes estimated to populate the entire intestine [52], and strongly depending on growth and mixing characteristics (**Fig. 4C**). Notably, this number is an upper bound for the effective population size of specific species as a single species account for only a subfraction of all microbial biomass. For example, if a microbial species accounts for 1% of all microbial cells, the effective population size varies between 10^7^ and 10^9^ cells. The emptying of the cecal compartment might also be more actively promoted in contrast to what is modeled here, lowering 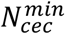 even further. Overall, these simulation results thus highlight how populations undergo strong bottlenecks several times a day with possibly strong effects on the evolution of the gut microbiome.

**Figure 4.**
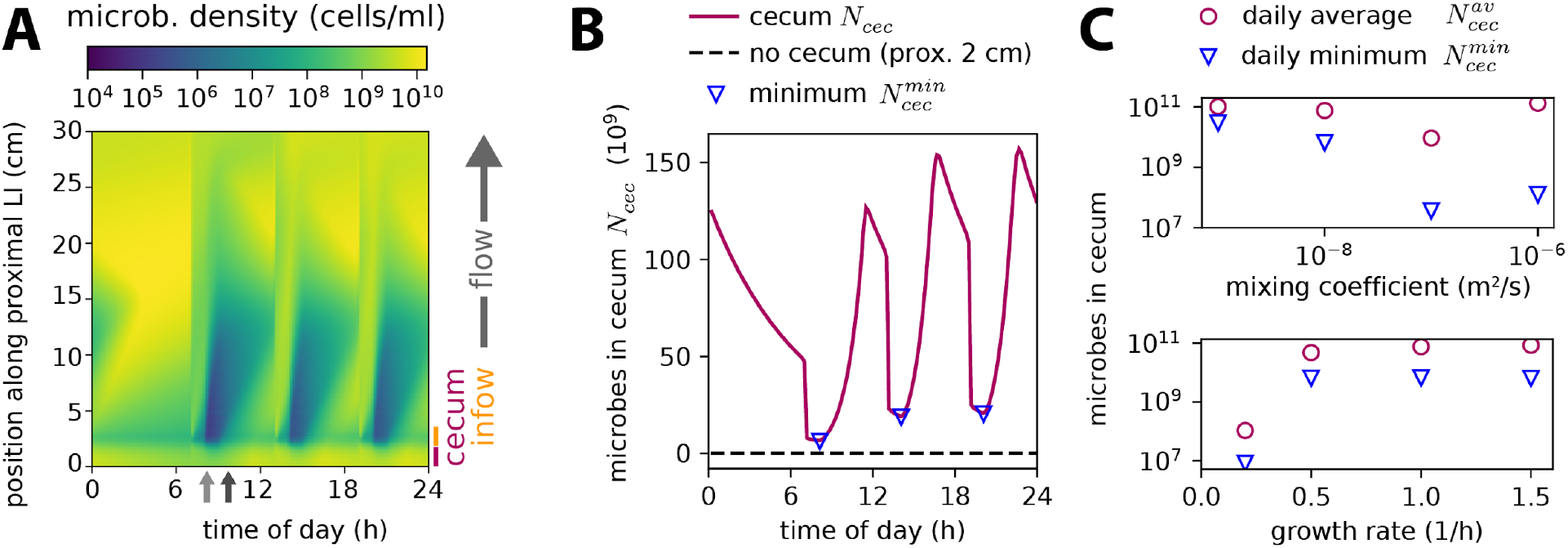
The microbial population in the cecum. **(A)** Kymograph showing radially averaged microbial density for the full model scenario with extendable intestinal volume and a cecal compartment. After meal intake, peaks in fluid supply lead to a rapid decrease in microbial population densities along the ascending colon (gray arrow). The cecal population then seeds growth (black arrow) and thus sets the effective population size (see text). **(B)** Temporal variation in total number of cecal microbes (magenta, *N*_*cec*_). The bottleneck size (triangular markers), shown as minimum population 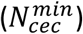, determines the effective population size of the system. For the case with “no cecum”, numbers in the proximal 2 cm of the LI are shown. **(C)** Daily average (magenta, 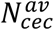) and minimum (blue, 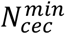)of the cecal microbial population versus variations in mixing conditions (top graph) and microbial growth rates (bottom graph). A growth rate of *λ* =1 1/h and mixing strength of *D* = 10^−8^ m^2^/s are used in the simulations for the top and bottom graphs, respectively. Other used parameter values are listed in **Table S1**.

### Variations in Growth Lead to Variations in the Uptake of Fermentation Products

To further investigate the consequences of diurnal growth variations for the host, we finally model the fermentation products (FPs) released by primary fermenters to grow within the anoxic environment of the LI. Given the relatively low energy yield of fermentation, microbes release high amounts of FPs. For example, up to about 90% of carbon in complex carbohydrates microbes consume is commonly released in the form of FPs [9]. Most of the FPs are taken up by the intestinal epithelium providing a substantial energy source for the host, 2-12% of the daily energy demand depending on diet and impacting many physiological processes [53,54]. We simulate this turnover of FPs by incorporating microbial excretion and host uptake into the modeling framework (**Fig. 5A** and **SI Text 3**). Rates describing the microbial release of FPs, *ε*_*FP*_, and the subsequent uptake per epithelium area, *J*_*FP*_, are taken from culturing experiments [9] and physiological studies of the human LI [55,56], respectively. Coupled with microbial growth dynamics, the concentration of FPs in the proximal LI varies significantly throughout the day, with particularly high concentrations during nighttime and lower concentrations during daytime (**Fig. 5B**, orange lines). Consequently, the uptake of FPs by the host also fluctuates strongly (**Fig. 5B**, blue line), leading to a substantial decrease in FPs concentration within the intestine (**Fig. 5B**, grey lines). Notably, this uptake results in the gradual decrease of FPs across the cross-section of the LI, particularly at the intestinal wall (**Fig. 5C**), which also impacts host uptake. Thus, the uptake rate does not only depend on the epithelial uptake capacity, *J*_*FP,max*_, but also on the mixing strength, *D*, of luminal contents (**Fig. 5D**).

**Figure 5.**
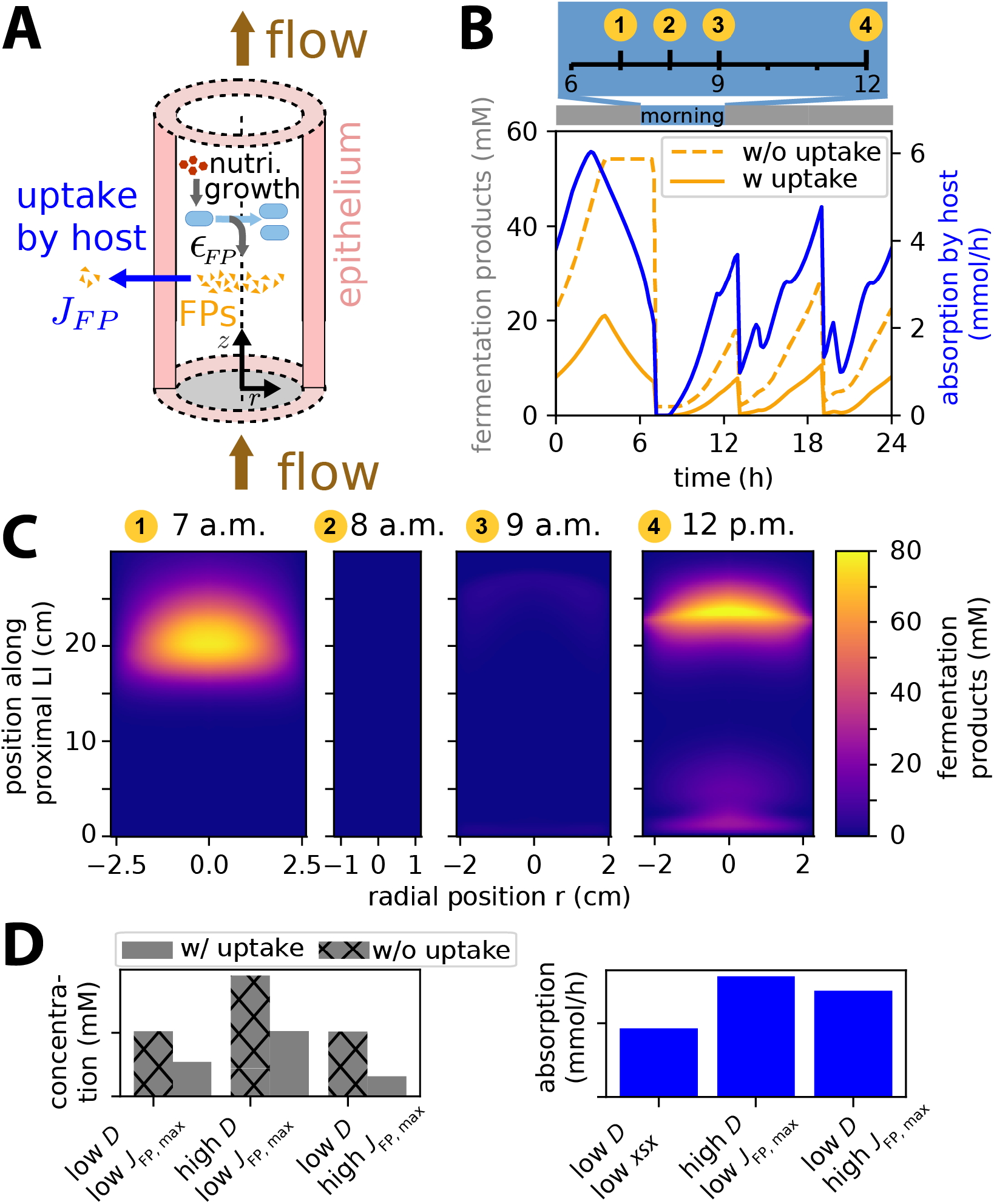
Spatiotemporal variation of microbial FPs. **(A)** Model incorporating microbial secretion and subsequent uptake of FPs by the host. **(B)** Temporal variation in concentration (orange) and uptake (blue) of FPs along the proximal LI. **(C)** Spatial distribution of FPs concentrations at different times of the morning. Distribution of nutrients and microbes for the corresponding time points are shown in **Fig. 2D-E. (D)** Daily average of concentration and uptake for three different conditions with low and high mixing strengths, *D* = 10^−8^ and 10^−6^ *m*^2^*/s*, and low and high maximum uptake rates, *J*_*FP,max*_ = 2.5 × 10^−5^ and 7.5 × 10^−5^ *mol/*(*m*^2^ · *s*). Additional parameter values are listed in **Table S1**. Time-lapsed dynamics are shown in **Video S1**.

## Discussion

In this study, we have examined how strong diurnal variations in nutrient supply and luminal flow shape microbial population dynamics along the proximal LI. Through the integrated consideration of flow, microbial growth, cycles of volume expansion/contraction, and mass movements, our simulation results emphasize significant fluctuations in microbial population sizes throughout the day. Crucially, changes in the intestinal volume and the presence of a dead-end cecal compartment off major flow paths are essential in ensuring the stable microbial density observed in the proximal LI. Microbial growth rates and the mixing of luminal contents further impact the dynamics, which include strong population bottlenecks and the recovery of microbial populations after large inflow events following meal intakes.

These findings reveal strong consequences on the microbiota and its interaction with the host. For example, the intertwined dynamics discussed here have strong consequences on evolution simply because of changes in population size: Caused by digestion-dependent flow control and nutrient supply, the population cycles through bottlenecks with 10^7^ to 10^10^ cells several times a day. Different theoretical and in-vitro studies have shown that such cycling can strongly shape evolutionary dynamics [51,57,58]. Consider the long-term laboratory evolution experiment by Lenski and colleagues with the model gut bacterium *Escherichia coli* as illustration [59]. Daily dilution followed by growth leads to the cycling of population sizes with a bottleneck size of 10^7^, comparable to the population size bottlenecks in the gut and with well-established consequences of these bottlenecks on evolution [51,60].

Another important consequence is the limitation of cross-feeding on lower trophic levels: To metabolically utilize microbial excretion products in the anoxic environment of the gut, cross-feeding species need to employ metabolic strategies alternative to fermentation, such as methanogenesis or chain elongation [47]. Those strategies are metabolic inefficient, resulting in substantially slower growth compared to primary fermenters [48]. Given the strong flow, these slow-growing cross-feeders can thus not accumulate efficiently along the proximal large intestine, and microbial metabolites are instead mostly taken up by the host to serve as energy sources and signaling factors [54]. Furthermore, as the microbial secretion of FPs varies with nutrient availability throughout the day, their uptake by the host can also vary strongly, with possible impacts on host physiology and behavior, such as hunger and satiety control [61].

Finally, the simulation results emphasize the host’s various abilities to control and adjust microbial population dynamics, for example, via the regulation of luminal water turnover, the adjustment of mass movements, and the mixing of luminal contents. As such, our analysis underscores highly intertwined population dynamics, essential to consider for gaining a more mechanistic understanding of microbe-host interactions.

We based our modeling framework on the current physiological understanding of the large intestine, focusing on the most important processes and providing new insights into the functioning of the human microbiota not possible with other approaches, like studies with animal models. But naturally, this modeling approach comes with limitations as it simplifies many complex host processes. Future in-vitro experiments, in-vivo studies, and extended modeling are thus needed to probe model predictions and better elucidate the discussed intertwined dynamics and their feedback with other intestinal processes. For example, the release of FPs at high levels is known to acidify the luminal environment, which can differentially slow down the growth of various species, linking microbiota composition to nutrient intake and flow [26,62]. The diurnal variations in nutrient availability, microbial growth, and FPs turnover discussed in this study are expected to shape this interplay, with strong consequences for microbiota composition. Similarly, the differential utilization of nutrients might lead to a substantial variation in growth behavior, which, given the close connection between growth and population dynamics, shapes not only composition but also the absolute numbers of microbial cells. Finally, we think the role of the vermiform appendix should be further explored. It has been previously suggested to serve as a safehouse for promoting a stable recovery of the microbiota after drastic disruptions, such as diarrhea or antibiotics treatment [63–65]. Given the regular cycling of the microbiota through relatively small population sizes discussed in this study, the presence of a substantial microbial population in the appendix [63,66], and the observed changes in microbiota composition after appendectomy [65], future investigations might reveal a more continuous role of the vermiform appendix in controlling and stabilizing the gut microbiota on a daily basis.

## Author Contributions

A.S. and J.C. developed the research project, analyzed and discussed the data, and wrote the manuscript. A.S. implemented and ran the finite-element simulations.

## Acknowledgment

We thank Markus Arnoldini, Benjamin Good, and members of the Cremer group for critical reading and suggestions to improve the manuscript.

## Code and Data Availability

A simulation file to run COMSOL Multiphysics simulations, including different elements of the model, is available on the GitHub repository accessible via the website https://github.com/cremerlab/diurnal_growthvariations. This repository also contains the Python code to analyze simulation results and generate figures.

