## Supplementary Tables, Text, and Figures for "Diurnal variations in digestion and luminal flow determine microbial population dynamics along the human large intestine"

### Supplementary Information

Alinaghi Salari[1,2,3] and Jonas Cremer[3*]

[1] Department of Chemistry, University of Toronto, Toronto, ON M5S 3H6, Canada
[2] Terrence Donnelly Centre for Cellular & Biomolecular Research, University of Toronto, Toronto, ON M5S 3E1, Canada
[3] Department of Biology, Stanford University, Stanford, CA 94305, USA

### Supplementary tables

**Table S1.** List of the variables/parameters used in this study.

| Parameter | Value | Description |
|---|---|---|
| $c_n$ | Variable | Concentration of nutrients |
| $c_b$ | Variable | Concentration of microbes (bacteria) |
| $c_{FP}$ | Variable | Concentration of FPs |
| $Y$ | 0.094 | Microbial growth yield in OD600 per mM of glucose (typical value for Firmicutes) [1] |
| $\lambda$ | 0.2, 0.5, 1, and 1.5 $(1/h)$ | Maximum growth rate of microbes (commonly observed in in-vitro studies) |
| $K$ | 50 $(\mu M)$ | Monod constant [1] |
| $K_{FP}$ | 20 $(\mu M)$ | Uptake constant for FPs [1] |
| $\varepsilon_{FP}$ | 12.5 | Excretion of FPs in mM per OD600 of microbes [1,2] |
| $J_{FP,max}$ | $2.5 \times 10^{-5}$ and $7.5 \times 10^{-5}$ $(mol/(m^2 \cdot s))$ | Maximum uptake rate of FPs by the epithelium |
| $D$ | $10^{-6}, 10^{-7}, 10^{-8}$, and $10^{-9}$ $(m^2/s)$ | Diffusion coefficient (See **SI Text 4**) |
| $\mu$ | $10^{-3}$ (Pa.s) | Dynamic viscosity of the colonic fluid (similar to water) |
| $\rho$ | $10^3$ $(kg/m^3)$ | Density of the colonic fluid (similar to water) |
| $p$ | $10^5$ (Pa) | Pressure |
| $L$ | 30 $(cm)$ | Length of proximal LI simulated [1] |
| $R_1$ | 0.97 $(cm)$ | Initial radius of proximal LI [1] |
| $R_2$ | Variable | Radius of proximal LI at the start of the mass movement |
| $R$ | $\sim 0.97 (cm) \leq R \leq \sim 2.7$ $(cm)$ | Radius of the proximal LI at any time point (corresponding to a volume range between 0.2 and 0.9 L) |
| $d_{ileo}$ | 2 $(cm)$ | Diameter of ileocecal opening [3] |
| $L_{ileo}$ | 5.16 $(mm)$ | Adjusted length of the ileocecal valve (Eq. [S16]) |
| $b_i$ | 5 (OD600) | Initial density of microbes |
| $c_{n_{in}}$ | See **Fig. S5** | Nutrient concentration |
| $c_{n_{in,initial}}$ | 0 | Initial concentration of nutrient |
| $c_{n_{in,basal}}$ | 0 | Basal (off-peak) concentration of nutrient at the inlet |
| $Q_{in}$ | See **Fig. S5** | Flow rate at the inlet for the cases of varying inflow assumption |
| $Q_{in,constant}$ | 62.5 (mL/h) | Flow rate at the inlet for the cases of constant inflow assumption |
| $Q_{in,basal}$ | 29.4 (mL/h) | Basal (off-peak) flow rate at the inlet for the cases of varying inflow assumption |
| $u$ | Variable | Velocity field with the components of $u_r$, $u_z$, and $u_\theta$ |
| $u_{initial}$ | 0 | Initial velocity field |
| $u_{out}$ | 0.7 (cm/s) | The emptying velocity because of contractions of the proximal LI |
| $t_m$ | Variable | Duration of mass movement |

| | | |
|---|---|---|
| $p$ | Variable | Pressure |
| $r, z, \theta, t$ | - | Spatial coordinates and time |
| $M_{LI}$ | See **SI Text 1** | Total microbial biomass in the LI |
| $D_L$ | See **SI Text 1** | Microbial loss rate from the LI |
| $T_t$ | See **SI Text 1** | Transit time through the intestinal tract |
| $V_{LI}$ | See **SI Text 2** | Volume of the LI |
| $L_{LI}$ | See **SI Text 2** | Length of the LI |
| $Q_{LI}$ | See **SI Text 2** | Average flow rate through the LI |
| $A_{LI}$ | See **SI Text 2** | Cross section area of the LI |
| $\phi$ | See **SI Text 2** | Turnover rate of microbes |
| $t_{dbl}$ | See **SI Text 2** | Doubling time of microbes |
| $t_{lum}$ | See **SI Text 2** | Time microbes remaining in a bioreactor/intestine |
| $N$ | See **SI Text 2** | Number of bioreactors |
| $t_{mix}$ | See **SI Text 2** | Mixing time scale |
| $t_{adv}$ | See **SI Text 2** | Advection time scale |
| $t_{transport}$ | See **SI Text 2** | Transport (combined mixing and advection) time scale |
| $l_{mix}$ | See **SI Text 2** | Mixing length scale |
| $l_{adv}$ | See **SI Text 2** | Advection length scale |
| $t_{growth}$ | See **SI Text 2** | Growth length scale |
| $N_e$ | Variable | Effective population size in the proximal LI |
| $N_{cec}$ | Variable | Population size in the cecum |
| $N_{cec}^{min}$ | Variable | Daily minimum population size in the cecum |
| $N_{cec}^{av}$ | Variable | Daily average population size in the cecum |

### Supplementary figures

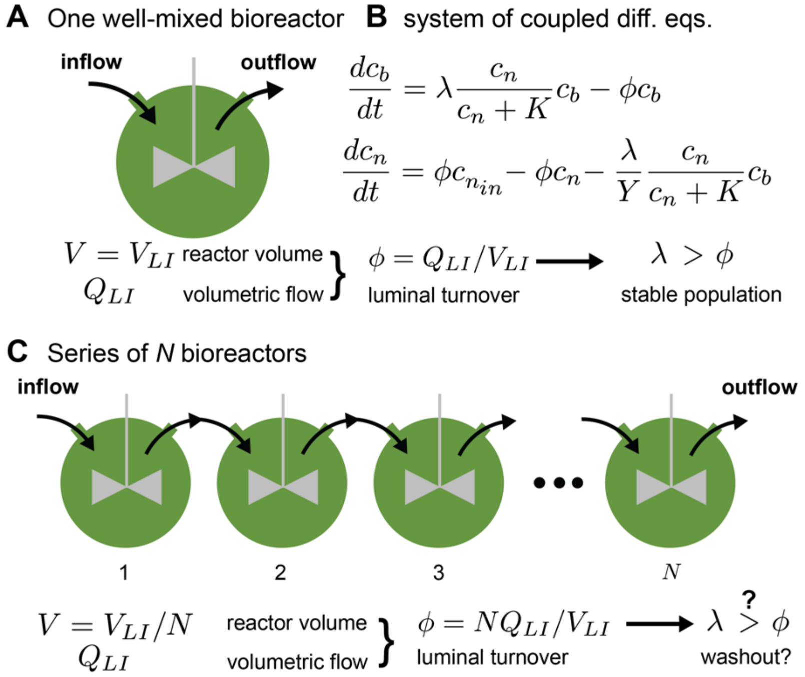

**Figure S1. Well-mixed bioreactor models.** Modeling microbial growth and nutrient consumption in the intestine by one or a series of well-mixed bioreactors. These models illustrate the fundamental link between growth, flow, and nutrient turnover. **(A, B)** Setup and equations for a single well-mixed bioreactor. For a stable population to be maintained, growth rate must exceed the turnover of luminal contents which is given by the ratio of the flow rate of luminal contents through the reactor and the reactor volume. **(C)** An array of coupled equally-sized bioreactors. The same equations describe growth and nutrient consumption in each reactor. However, because of a reduction of each reactor volume, turnover rates are higher, and the washout condition depends on the number of reactors, *N*. See **SI Text 2** for detailed introduction and discussion.

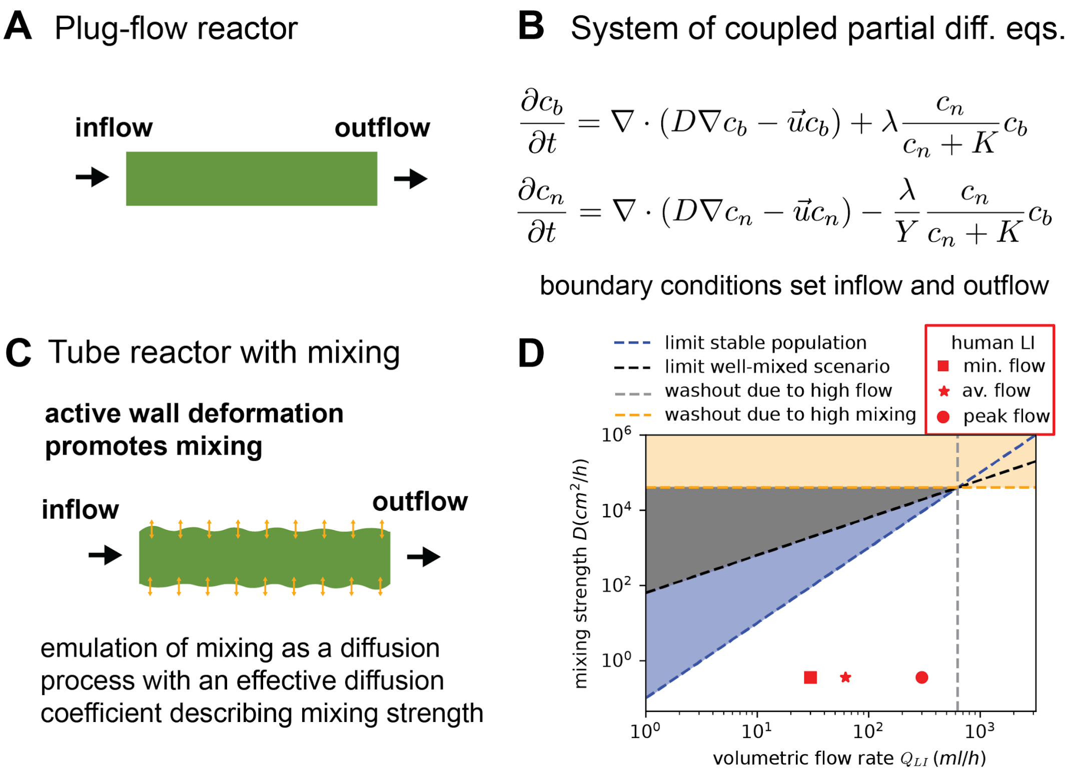

**Figure S2. Continuous tube models.** Modeling growth and nutrient consumption in a continuous tube. **(A, B)** Setup and 1-dimensional equations to describe nutrient consumption and microbial growth in a plug-flow reactor. **(C)** Tube reactor setup with an active mixing component. This is the same setup as a plug-flow reactor, but it accounts for active mixing via a diffusion process with a (relatively high) diffusion coefficient setting the strength of mixing. **(D)** Phase diagram illustrating regimes of stable population growth and washout in this setup, depending on volumetric flow rate $Q_{LI}$ and mixing strength $D$. See text for description of scaling relations. A growth rate of $\lambda = 1\ 1/h$ and tube cross-section area of $A_{LI} = 3.14\ cm^2$ are assumed. Square, star, and circle markers represent the minimum, average, and maximum flow rates, respectively, that are used in our simulations (See **Fig. S5**) at a mixing strength of $D = 10^{-8}\ m^2/s$. These findings suggest that an additional process beyond mixing is needed to ensure stable microbial densities in the gut despite the presence of strong flow. See **SI Text 1** for a detailed introduction and discussion.

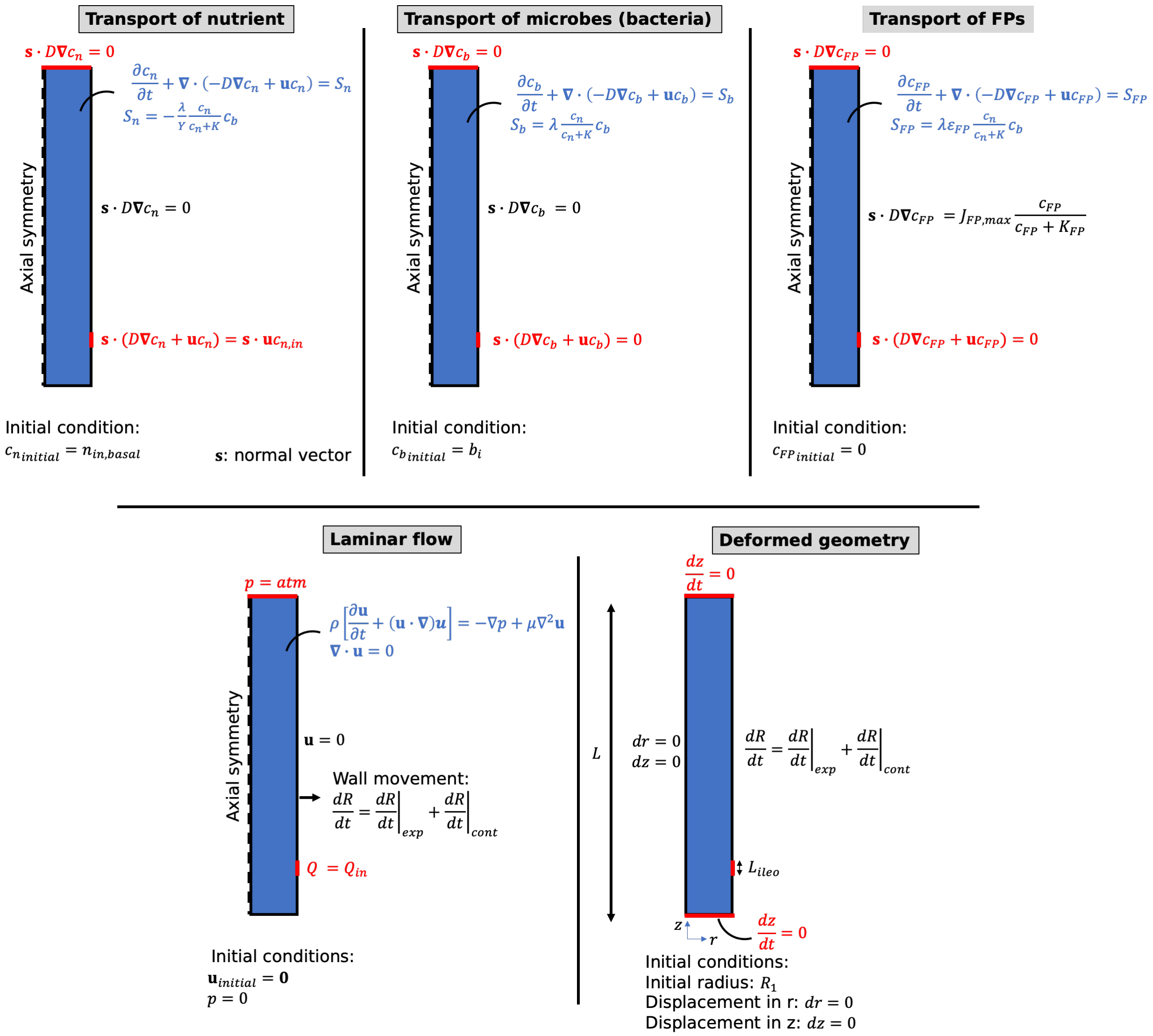

**Figure S3. Boundary and initial conditions**. Setups for different quantities and physical properties, labeled as transport of nutrients, transport of microbes (bacteria), transport of FPs, laminar flow, and deformed geometry, for the 2D axisymmetric simulation domain. The inlet and outlet of the domain, as well as their corresponding boundary conditions, are highlighted in red color. The simulation domain and the system of equations solved for it are highlighted in blue color. The boundary conditions applied to the intestinal walls are depicted in black color. With the axisymmetric setup of the simulation domain, the ileocecal valve has to be defined as a ring with a perimeter equal to the perimeter of the intestine and an adjusted width of $L_{ileo}$ (See **Table S1**).

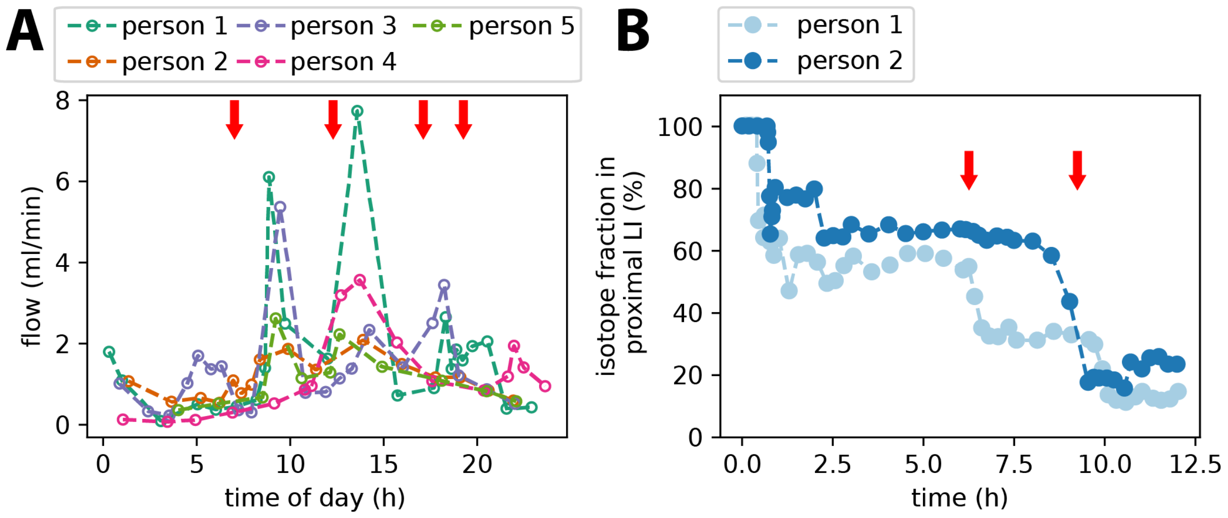

**Figure S4. Experimental observations confirm mass movements and temporal variation in inflow coupled with meal intake. (A)** Luminal water inflow reported by Phillips and Giller [4]. In their study, the dilution of infused polyethylene glycol solutions was used to calculate the flow through the ileocecal valve for different healthy subjects consuming three major and one smaller meal throughout the day (red arrows). Notably, except for one individual not showing a peak after breakfast and two not showing peaks after supper, the inflow shows strong variations with peaks in inflow occurring after meal intake. The limited time resolution (markers) thus already establishes the existence of inflow peaks, but inflow peaks might be even more pronounced in the real gut and better observable if data with higher time resolution were available. **(B)** Emptying of the ascending colon over time was analyzed by Hammer and Phillips [5]. In their study, 100 mL of radiolabelled isotopes were infused into the cecum of healthy subjects at a constant rate over a short period of 30 min. The isotope signal was then tracked over hours while subjects continued taking meals (red arrows). Notably, the decrease in signal occurs in a non-linear way with drops coordinated with meal intakes and indicating the rapid emptying of the ascending colon after meal intake.

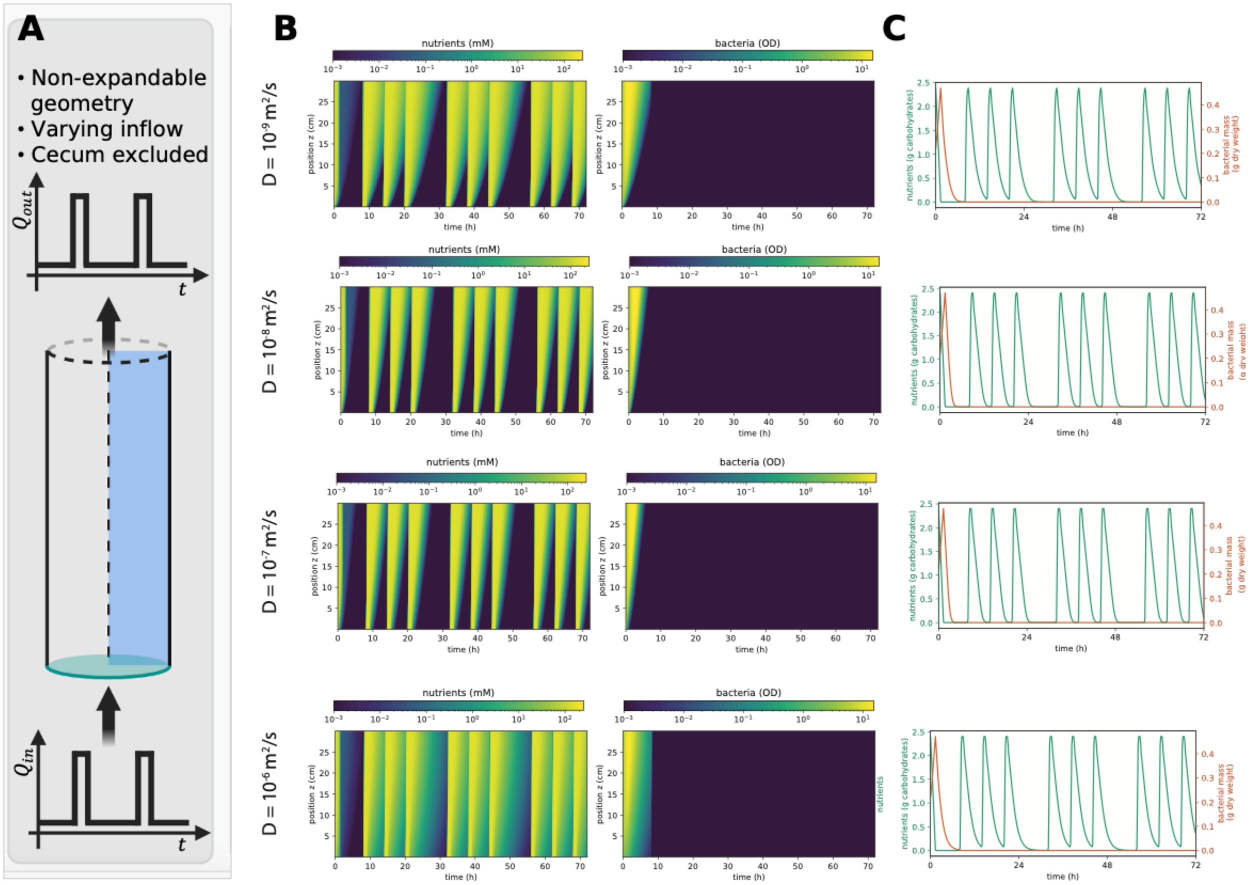

**Figure S5. Results for the case of varying inflow without cecum or mass movements.** Scenario as in **Fig. 1D**. **(A)** Schematics of setup. **(B, C)** Simulation results for four different mixing coefficients. **(B)** Kymographs showing the radially averaged nutrient concentrations and microbial densities over a 72-h simulation. **(C)** Total nutrients and microbes (bacteria) within the proximal LI versus the simulation time. In this scenario, no substantial bacterial population remains after ~10 hours. The system of equations and the list of parameters used in the simulations are shown in **SI Text 3** and **Table S1**, respectively. The inflow over time is shown in **Fig. 1C**.

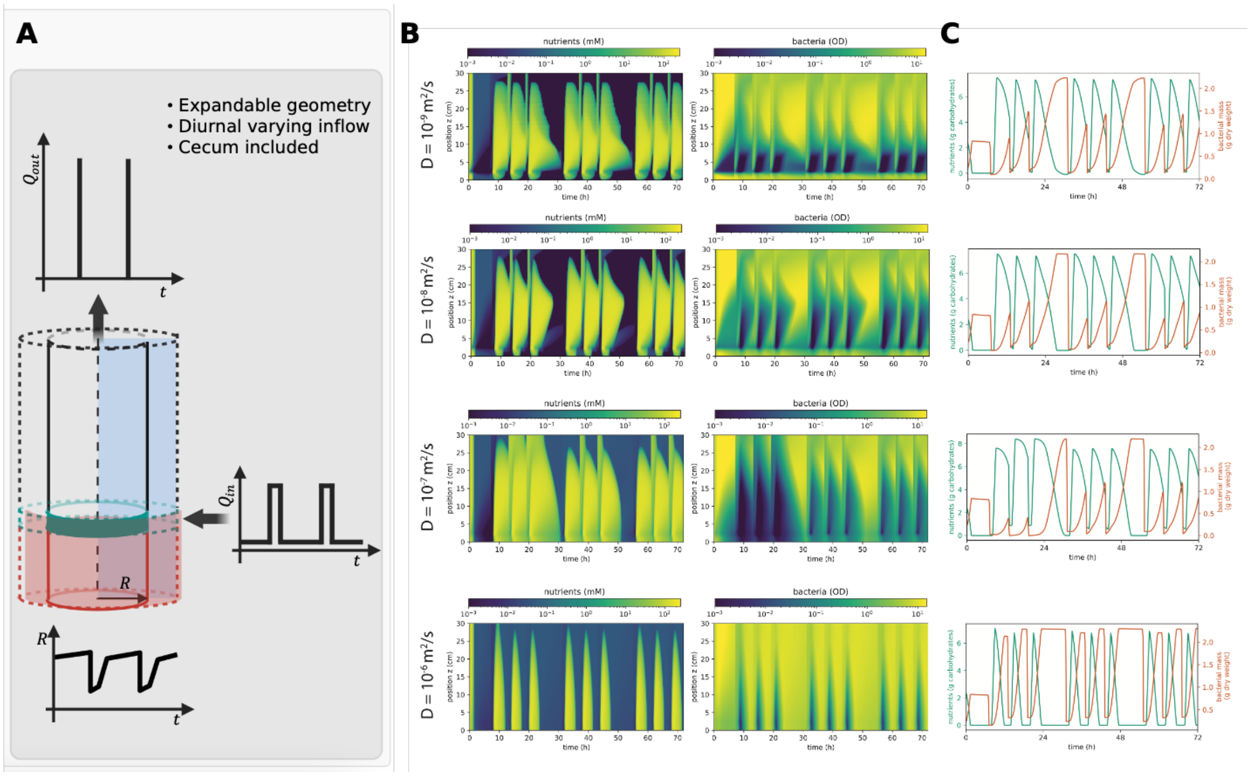

**Figure S6. Results for the case of varying inflow with cecum and mass movements included (full model scenario).** Scenario as in **Fig. 1F**. **(A)** Schematics of setup. **(B, C)** Simulation results for four different mixing coefficients. **(B)** Kymographs showing the radially averaged nutrient concentrations and microbial densities over a 72-h simulation. **(C)** Total nutrients and microbes (bacteria) within the proximal LI versus the simulation time. After the temporary variations over the first 24 h, the bacteria/nutrient profile follows a regularly fluctuating pattern with overall high bacterial densities. The system of equations and the list of parameters used in the simulations are shown in **SI Text 3** and **Table S1**, respectively. The inflow over time and the radius variation are shown in **Fig. 1C** and **2A**, respectively. As a result of mass movement, the outflow occurs in sharp peaks after meal intake.

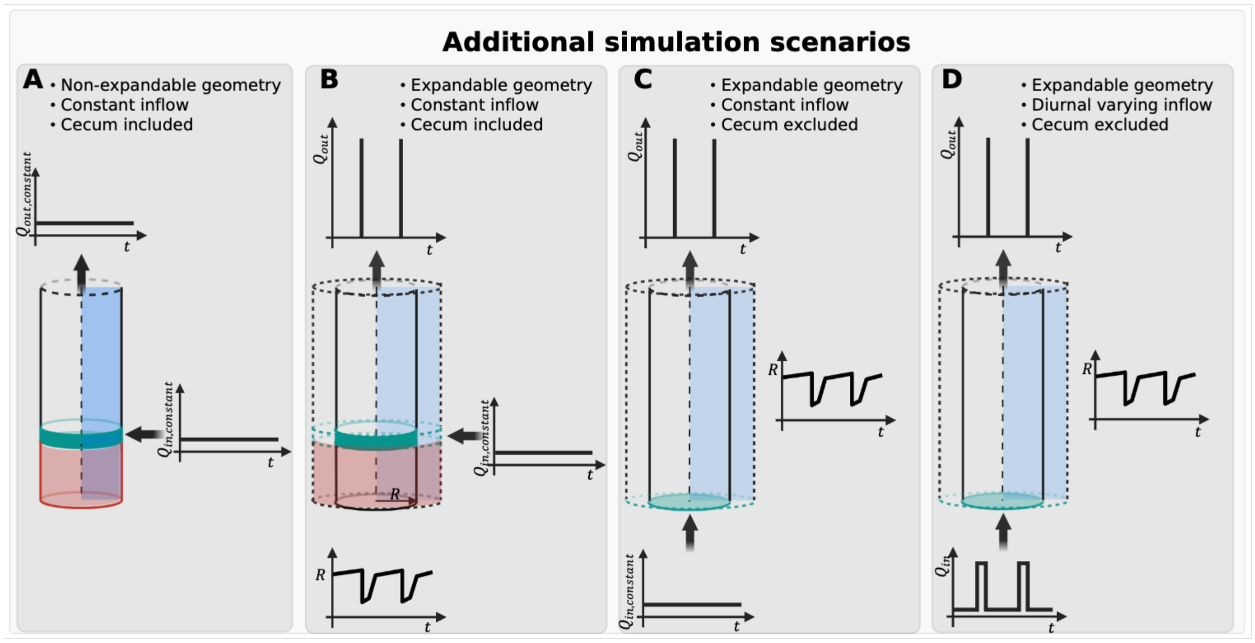

**Figure S7. Different non-physiological scenarios of flow and growth.** In addition to those scenarios discussed in the main text, we analyze more non-physiological scenarios with various inflow conditions, the presence or absence of a cecum, and the expandability of the proximal LI, for comparison. **(A)** Constant inflow, cecum, no radial expansion. **(B)** Constant inflow, cecum, radial expansion. **(C)** Constant inflow, no cecum, radial expansion. **(D)** Variable inflow, no cecum, radial expansion. The system of equations and the list of parameters used in the simulations are shown in **SI Text 3** and **Table S1**, respectively. The simulation results for these scenarios are shown in **Figs. S8 and S9**.

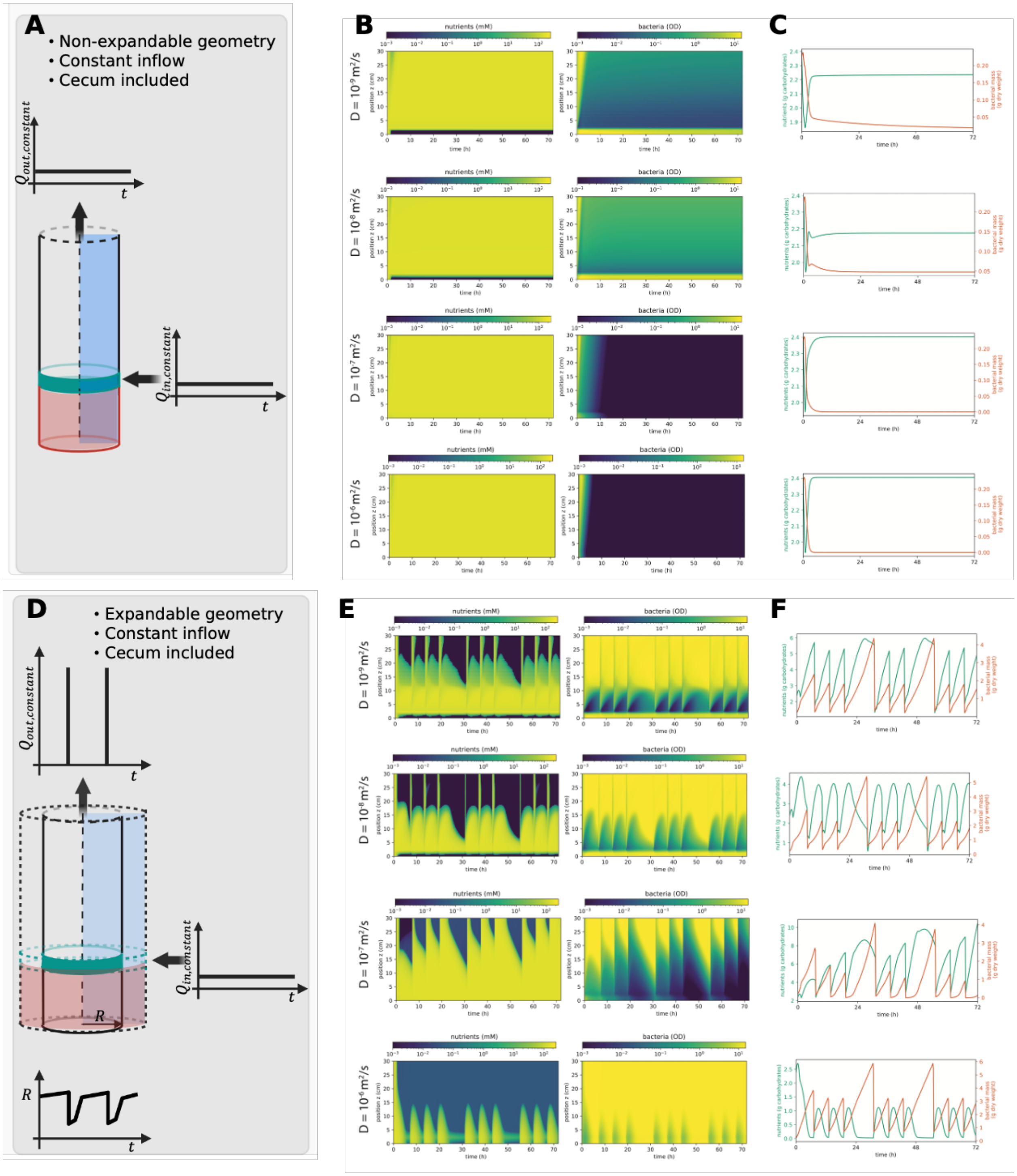

**Figure S8. Dynamics when cecum is present with constant inflow.** Simulation results when inflow is constant, and a cecum is present. **(A-C)** Scenario with a fixed radius, showing schematics (A), kymographs indicating nutrient and microbial density dynamics (B), and the total nutrients and microbes (bacteria) within the proximal LI over 72 h. The inflow and outflow are always constant and equal to $Q_{in,constant}$. **(D-F)** Scenario accounting for mass movement and a variable radius, showing schematics (A), kymographs indicating nutrient and microbial density dynamics (B), and the total nutrients and microbes within the proximal LI over 72 h. While the inflow is constant, the outflow fluctuates with sharp peaks post meal intakes related to mass movements. Kymographs show radially averaged nutrient concentrations and microbial densities. The system of equations and the list of parameters used in the simulations are shown in **SI Text 3** and **Table S1**, respectively. The radius variation is shown in **Fig. 2A**.

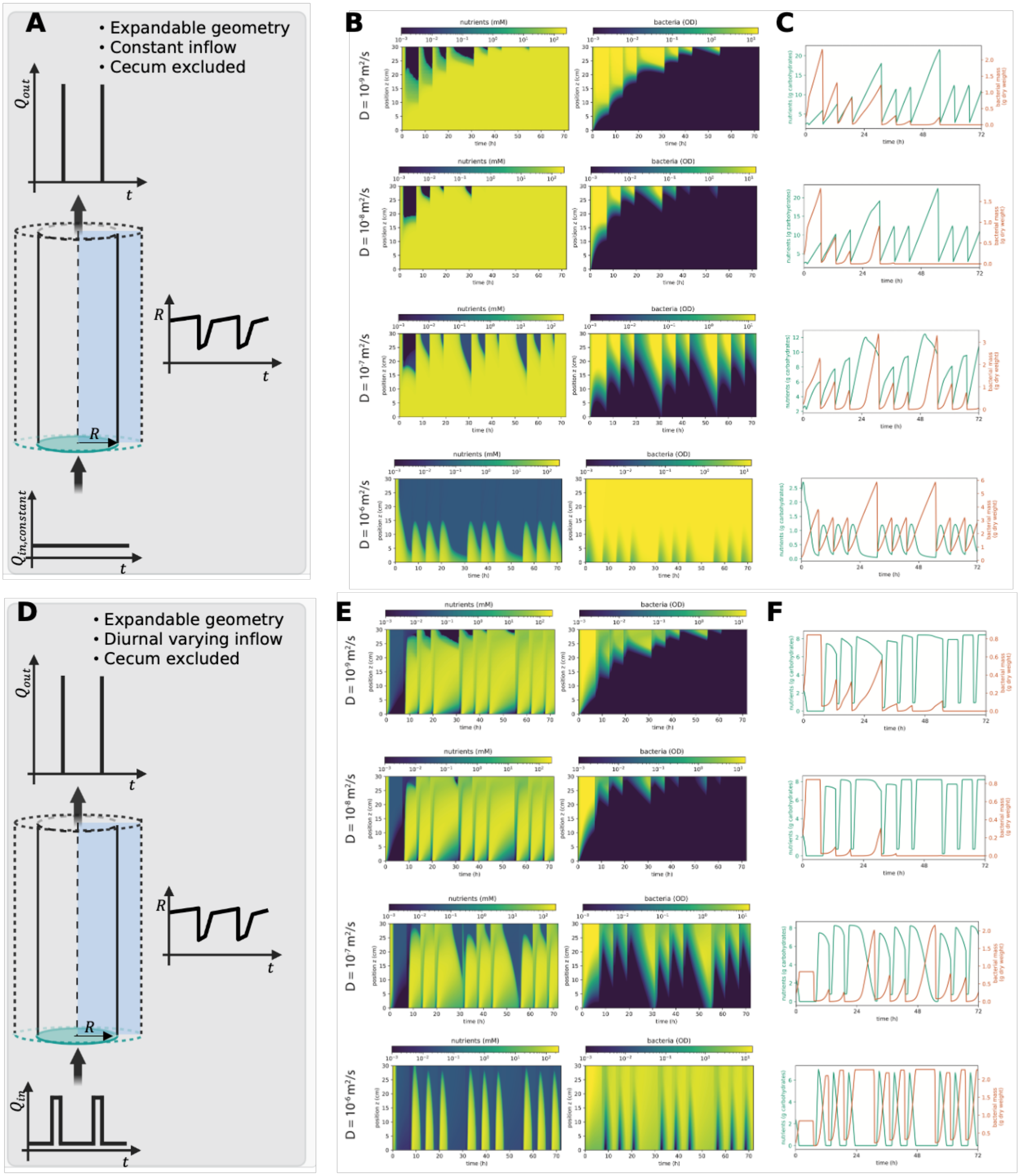

**Figure S9. Dynamics without cecum but variable colonic radius.** Simulation results when no cecum is present, but radius is variable and controlled by mass-movements. **(A-C)** Scenario with a fixed inflow, showing schematics (A), kymographs indicating nutrient and microbial density dynamics (B), and the total nutrients and microbes (bacteria) within the proximal LI over 72 h. While the inflow is constant, the outflow fluctuates with sharp peaks post meal intakes related to mass movements. **(D-F)** Scenario accounting for inflow changing over time, showing schematics (A), kymographs indicating nutrient and microbial density dynamics (B), and the total nutrients and microbes within the proximal LI over 72 h. Both the inflow and the outflow fluctuate with sharp peaks in meal intakes related to digestion and mass movements. Kymographs show radially averaged nutrient concentrations and microbial densities. The system of equations and the list of parameters used in the simulations are shown in **SI Text 3** and **Table S1**, respectively. The radius variation is shown in **Fig. 2A**.

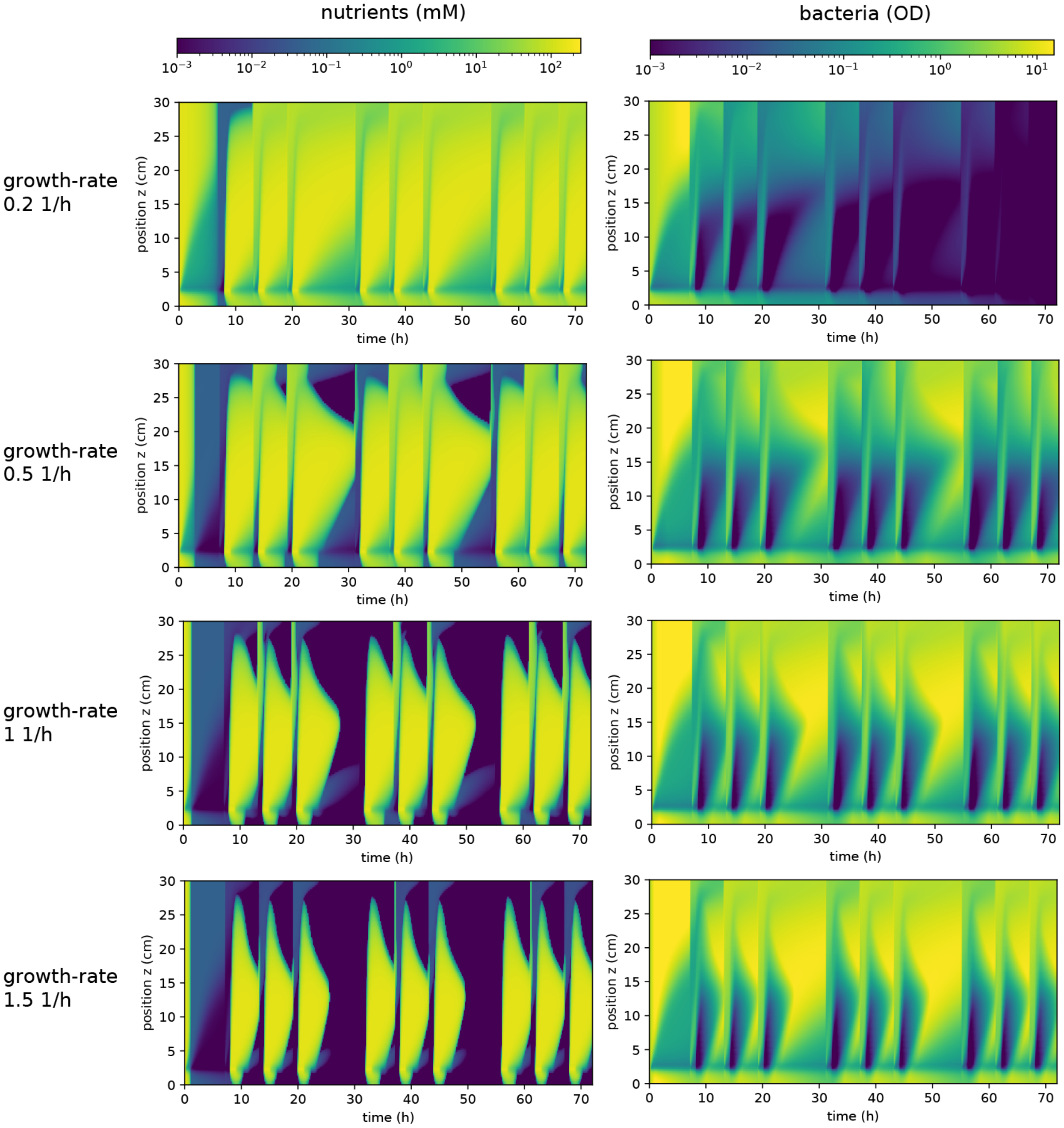

**Figure S10. Variation of growth rate for the full model scenario.** Simulation results for the case of varying inflow, cecum, and mass movements included (full model scenario, **Fig. 1F**) for four different growth rates. Kymographs show the radially averaged nutrient concentrations and microbial densities over a 72-h simulation. For slow growth (growth rates of 0.2 1/h), the microbial population gradually decreases over time, leading to a washout over the simulated time. In contrast, a stable population remains for higher growth rates. The system of equations and the list of parameters used in the simulations are shown in **SI Text 3** and **Table S1**, respectively.

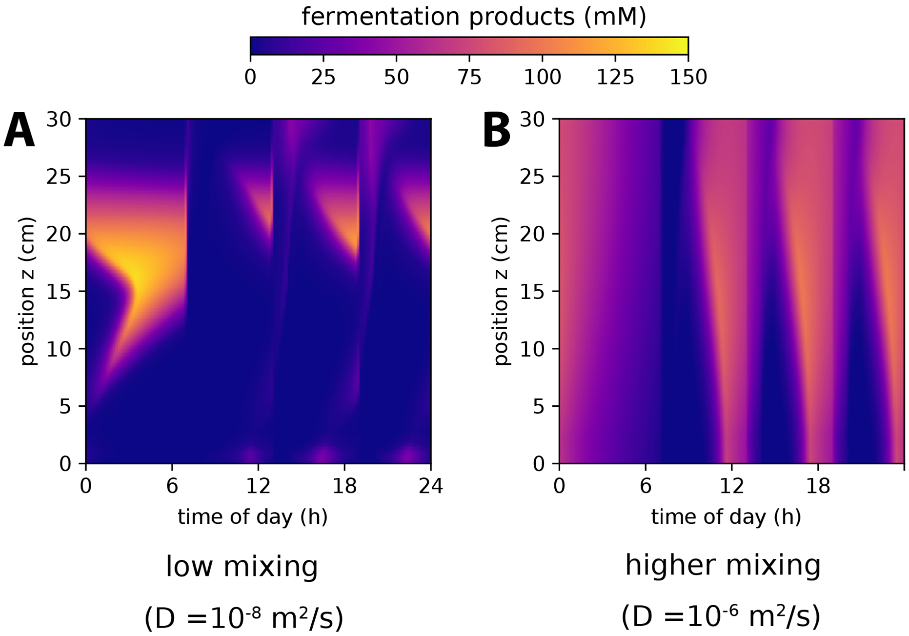

**Figure S11. Spatiotemporal variation of FPs. (A, B)** Kymographs show spatially averaged concentration of FPs along the LI and throughout the day, for lower (A) and higher (B) mixing strengths, as indicated. Simulation parameters are the same as those used in **Fig. 5** with FPs uptake rate of $J_{FP,max} = 2.5 \times 10^{-5}\ mol/(m^2 \cdot s)$.

### Supplementary videos – captions

**Video S1. Temporal variation of nutrients, microbial densities, and fermentation products.** The video shows the variation of nutrient concentration (upper row), microbial density (center row), and FPs (lower row) throughout the day (time of day indicated) for the full model scenario (with mass-movements and cecum included). Plots on the left illustrate full spatial variation of the three quantities. Plots on the right show kymographs of the radially averaged quantities, with a moving white dashed line indicating the timepoint for the spatial variation shown on the left. The same parameters as those used in **Figs. 2 and 5**, with parameters listed in **Table S1**, including a mixing strength of $D = 10^{-8}\ m^2/s$ and a FPs uptake rate of $J_{FP,max} = 2.5 \times 10^{-5}\ \frac{mol}{m^2 \cdot s}$, are used in this simulation.

### Supplementary text

### 1. Average growth and biomass turnover in the human large intestine

With a negligible number of microbes entering the large intestine (LI) from the small intestine [6], the total microbial biomass in the LI, $M_{LI}$, is the result of microbial growth minus their loss via feces. Mathematically,

$$\frac{dM_{LI}}{dt} = \lambda M_{LI} - D_L M_{LI} \quad \text{[S1]}$$

Here, $\lambda$ denotes the (spatially averaged) growth rate and $D_L$ denotes the loss rate (in bioreactor theory often called dilution rate). To maintain a stable microbial population, the time-averaged growth and loss rates need to match, i.e., $\langle\lambda\rangle = \langle D_L\rangle$. To estimate the average loss rate and, thus, the average growth rate, we can consider the time needed for contents to pass the intestinal tract. This transit time, $T_t$, is mostly determined by the transit through the LI. The loss rate can thus be estimated as $\langle D_L\rangle \approx ln\ ln\ (2)\ /T_t$. Notably, the transit time varies broadly with diet and the turnover of intestinal water along the intestine, ranging between 15 and 100 h [2,7]. Consequently, the average growth rate $\langle\lambda\rangle$ needs to fall within the range of 0.007 1/h and 0.05 1/h for a stable microbial population to form.

Importantly, this estimation provides a time- and space-averaged growth rate. As, for example, indicated by spatial variations in the concentration of fermentation products (FPs), in-vitro growth measurements of gut strains, or diurnal variation of nutrient supply considered in this study, actual growth rates can vary drastically in space and time. Particularly, growth rates may range from no growth to much higher maximal growth rates when local growth conditions are favorable, and nutrients are abundant.

### 2. Mathematical models for analyzing microbial growth and biomass turnover within the large intestine

Mathematical models of gut microbiota growth commonly build on bioreactor theory [8–13]. In this section, we introduce different models and their limitations for modeling microbiota in the LI. These considerations (i) illustrate the fundamental relation between growth, flow, and biomass turnover and (ii) clarify the relations of previous models to the modeling framework we introduce in this study.

#### 2.1. Scenario 1 – One well-mixed bioreactor

In the simplest case, fermentation is considered in a well-mixed bioreactor with continuous flow (**Fig. S1A**). Nutrient-enriched fluids continuously enter via an inlet, and biomass, as well as non-consumed nutrients, exist via an outlet. As we explain and emphasize below, such a model cannot adequately represent microbial (bacterial) growth dynamics in the LI. However, it well illustrates the fundamental balance between growth and fluid turnover in controlling microbial abundance.

Mathematically, we can describe growth, nutrient consumption, and fluid turnover in this scenario with two differential equations (**Fig. S1B**). The two most important parameters are the maximum growth rate, $\lambda$, which describes growth in nutrient-replete conditions, and the luminal turnover rate, $\phi$, which describes the rate with which luminal contents is replaced by new fluids (in bioreactor theory, this rate is commonly called the dilution rate). In a bioreactor with a fixed volume, $V_{LI}$, the turnover rate can be defined as the rate of volumetric flow passing through the LI, $Q_{LI}$, divided by its volume, i.e., $\phi = \frac{Q_{LI}}{V_{LI}}$. Stable growth is only ensured if this turnover (dilution) is lower than the maximal growth rate, i.e., $\phi < \lambda$. In other words, to ensure a stable biomass, the time microbes require to replicate, $t_{dbl} = \frac{ln ln\,(2)}{\lambda}$, needs to be shorter than the time luminal contents remain in the bioreactor, $t_{lum}$. To probe if this condition is fulfilled in the human LI, we can estimate the values for the growth and turnover rates.

1. First, measurements in nutrient-rich conditions have confirmed that most abundant gut bacteria commonly double every 45 minutes or faster, thus, $\lambda \geq 0.9\ 1/h$. [2]
2. Second, the adult human LI has a maximum internal volume of approximately $V_{LI} \approx 1$ L (upper bound, assuming ascending, transverse, and descending colon of $\sim$ 300 mL each) [14]. Typically, around 1.5 − 2 L of luminal contents enter the LI per day [15]. As there is substantial water uptake along the LI and not all water exits via feces, we consider this as an upper bound for the flow rate, $Q_{LI} = 2\ L/day \approx 83\ mL/h$. Using this, we get $\phi = \frac{83\ mL/h}{1000\ mL} \approx 0.08\ 1/h$ as an upper bound for the turnover rate in a well-mixed bioreactor.

Thus, the maximum luminal turnover rate appears to be substantially lower than the growth rate, $\phi \ll \lambda$, suggesting a stable microbial population can easily manifests itself within the LI. However, assuming the $\sim$ 2-m long LI operates like a well-mixed bioreactor is unrealistic. Instead, mixing is limited only to local intestinal regions, and thus this assumption drastically underestimates the conditions needed for a stable microbial population, as we consider next.

#### 2.2. Scenario 2 – An array of coupled bioreactors

Next, we consider the case of several coupled bioreactors, with the outflow of one bioreactor constituting the inflow of the next one (**Fig. S1C**). Such a setup is often realized in experimental studies with commonly 2 to 5 bioreactors being linked to emulate microbial growth in the intestine [8,12]. Importantly, for the consideration of stable microbial growth, the volume of each bioreactor in this model is smaller than the total volume of the intestine. If we assume $N$

equally sized reactors, each will have a volume of $V_{LI}/N$. However, the flow going through each bioreactor is still the same, approximately $Q_{LI} = 1.5\ L/day$ as discussed above. Therefore, the turnover rate is higher, i.e., $\phi = N \cdot 0.08\ h^{-1}$, and the requirement for a stable microbial population ($\phi < \lambda$) now depends on the number of bioreactors. As it is not directly apparent how many bioreactors are needed to emulate the growth dynamics along the human gut, it is not clear if gut conditions fulfill the requirement for stable growth. Or, phrased differently, the coupled bioreactor model can illustrate some basic characteristics of microbial growth and fluid turnover, but we need to go beyond this model to understand its limitations and the conditions that ensure a stable microbial population along the human LI.

#### 2.3. Scenario 3 - Continuous tube model with active mixing

To better understand flow conditions, we start with a simple continuous model that does not discretize the LI into different compartments but rather considers a continuous rigid tube. In bioreactor theory, such a scenario is known as plug flow reactor [16] (**Fig. S2A**). In the simplest form, a 1-dimensional tube can be assumed such that the transport equations accounting for the combined effects of nutrient consumption, microbial growth, and flow are coupled partial differential equations, as shown in **Fig. S2B**. Importantly, in this scenario, microbes are completely washed out of the tube even for the flow rates equal to the observed average gut flow rates [17]. Furthermore, the assumption of no active mixing of colonic contents is highly unrealistic, as there is ample evidence that the coordinated contraction of intestinal muscles gives rise to peristaltic movements. These movements not only promote the longitudinal transport of luminal contents but also enhance transverse mixing. A more recent study[8] has thus considered a model that also accounts for active mixing along the LI. This mixing is described by an effective diffusion term with a diffusion coefficient, $D$, as the mixing parameter, which is much higher than that of molecular diffusion (**Fig. S2C**). This approach was experimentally studied in vitro using a microfluidic channel that follows the dimensions of the mouse gut and allows for the emulation of active luminal mixing via the controlled active deformation of its channel walls.[18] Model predictions and microbial density measurements showed good agreement over a broad range of growth, flow, and mixing conditions, confirming this approach as a first-order approach to study mixing dynamics and their effect on growth.

Importantly, active mixing can promote a stable population as long as growth and mixing are fast compared to the volumetric flow rate. This is illustrated by a simple scaling argument and a phase diagram that states under which combinations of mixing and flow conditions one obtains a stable microbial population (**Fig. S2D**). The dynamics of this 1-dimensional system can be characterized using three timescales, each corresponding to one of the three governing phenomena, i.e., mixing (diffusion) $t_{mix}$, advection $t_{adv}$, and growth $t_{growth}$, as follows:

$$t_{mix} = \frac{l_{mix}^2}{D} \quad ; \quad t_{adv} = \frac{l_{adv}}{u_z} \quad ; \quad t_{growth} = \frac{1}{\lambda} \qquad \text{[S2]}$$

Here, $l_{mix}$ and $l_{adv}$ are mixing and advection length scales, respectively. By assuming advection and mixing having equal strength (i.e., *Peclect* number equal to one) in the transport of species along the tube, we can obtain a combined transport timescale as $t_{transport} = \frac{D}{u_z^2}$. For such a system to promote stable microbial densities, the timescale of growth should be smaller than the timescale of transport (mixing and advection combined), i.e., $t_{transport} > t_{growth}$. This means that $\lambda > \frac{u_z^2}{D}$ or $\lambda > \frac{Q_{LI}{}^2}{D.A_{LI}^2}$ or $D > \frac{1}{\lambda.A_{LI}^2} Q_{LI}{}^2$. Here, the axial velocity is defined as $u_z = \frac{Q_{LI}}{A_{LI}}$, where $Q_{LI}$ is the volumetric flow rate and $A_{LI}$ is the cross-sectional area. This condition is shown in **Fig. S2D** (in log-scale) as the region above the blue dashed line. The system furthermore reaches the limit of a well-mixed bioreactor if $l_{mix} = \frac{D}{u_z} > L_{LI}$, where $L_{LI}$ is the length of the LI, and thus $D > \frac{Q_{LI} \cdot L_{LI}}{A_{LI}}$ (area above the black dashed line). Under such well-mixed conditions, a stable population emerges if either $t_{mix}$ or $t_{adv}$ are larger than $t_{growth}$, which results in $\lambda > \frac{u_z}{L_{LI}} = \frac{Q_{LI}}{A_{LI} \cdot L_{LI}}$ (i.e., $\lambda. A_{LI}. L_{LI} > Q_{LI}$) (area to the left of the gray dashed line) or $\lambda. L_{LI}^2 > D$ (area below the yellow dashed line).

To approximate where on this phase diagram the LI operates, we estimate different parameters for the human gut. Assuming a mixing parameter of $D \approx 0.36\ cm^2/h$ (equivalent to $10^{-8}\ m^2/s$) and a volumetric flow rate of $83\ mL/h$ (as calculated above), the condition will be close to the washout limit (**Fig. S2D**, star). To address this unrealistic condition, a higher diffusion coefficient of $D = 36\ cm^2/h$ (equivalent to $10^{-6}\ m^2/s$) has been previously estimated [18] to be necessary for the emergence of a microbial population. For the case of higher diffusion coefficient, the mixing length scale is $l_{mix} = D \cdot \frac{A_{LI}}{Q_{LI}} \approx 1.4\ cm$, assuming a cross-section area of $A_{LI} = 3.14\ cm^2$. This suggests that a coupled bioreactor model containing about $N = \frac{L_{LI}}{l_{mix}} \approx 140$ bioreactors in series might be able to emulate the LI, regardless of the microbial growth rates. However, we stress here that mixing strongly varies with flow conditions which strongly change over time, suggesting that a coupled bioreactor setup with a fixed number of reactors is anyway problematic in recapitulating the growth dynamics along the LI.

In summary, the presented order of magnitude analysis illustrates how microbial growth and population densities depend on the subtle interplay between flow control, mixing, and growth. Minor variations in those aspects can potentially have very large effects on the stability and overall growth behavior of the microbial population. However, this analysis contains major simplifications which need to be probed further. For example, we only use the average volumetric flow rate, $Q_{LI}$, which, as discussed in the main manuscript, varies drastically throughout the day with distinct peaks in flow occurring after each meal intake when up to half a liter of fluids enters the LI within an hour ($Q_{LI} \approx 0.5\ L/h$ for the 1-h duration after meal intake; in our simulations we assume $0.3\ L/h$ during this 1-h). During these post-meal intake periods, the condition for a stable microbial population ($\lambda > \frac{Q_{LI}{}^2}{D.A_{LI}^2}$) is clearly not fulfilled, requiring unphysiological high growth rates of $\lambda \geq 2.5 \times 10^4\ 1/h$ when $D \approx 0.36\ cm^2/h$ and the flow rate is $0.3\ L/h$ (**Fig. S2D**, circle). This is further confirmed by our simulations which account for the observed variations in flow and model the full growth and nutrient consumption dynamics in a tube with active mixing: no stable microbial population remains even if mixing is very strong (See

**Fig. S5**). We think this is mostly because of another major simplification that the typical bioreactor models assume and which almost all in-vitro experiments incorporate, i.e., fixed volumes of the growth compartments that do not change over time. In contrast, as we emphasize in the main manuscript, the volume of the proximal LI is flexible. This flexibility reduces peaks in fluid turnover and, therefore, promotes a stable microbial population. With this dynamical change in volume, further processes need to be considered to gain a meaningful understanding of overall microbial growth, including the rapid emptying dynamics via mass movements or the consideration of the role of a cecum. Thus, we introduce and implement a more sophisticated model to capture these additional dynamics; see the following section for a step-by-step introduction

### 3. Formulation of the full mathematical model

#### Simulation domain

We begin the construction of our mathematical model by evaluating the volume of the proximal LI. Regarding the dimensions of our simulation domain, the measurements documented in the literature exhibit a broad range, depending on factors such as age, sex, and the methodology of measurement. The reported lengths for the cecum and ascending colon vary significantly, with values ranging from 2 to 14 cm for the cecum [19,20] and 9 to 62 cm for the ascending colon [1,4]. In our simulations, we assume that the ileocecal opening is positioned 2 cm from one end of the simulation domain. The cecum is considered to extend from the same end up to the ileocecal opening. The remaining ∼28 cm of the domain is designated as the proximal colon. We collectively refer to the entire simulation domain, measuring 30 cm in length and encompassing the cecum and proximal colon, as the proximal LI.

#### Volume change and mass movements

Based on data from the literature, the proximal LI can accommodate volumes ranging from less than 200 mL to about 900 mL [21–23]. In our full model, we assume that the proximal LI can expand to a maximum of 900 mL due to fluid intake. Notably, our simulations show that the volume of the simulation domain never reaches this maximum value before a mass movement is triggered, which pushes most colonic contents distally and out of the simulation domain, leading to a decrease in the simulation domain. Outside of mass movements, we assume that the proximal LI functions as an expandable container with an open inlet (i.e., the ileocecal opening) and no outflow. We do not impose a zero-outflow condition at the outlet. However, the actual outflow is zero because we set the rate of volume expansion equal to the rate of fluid entering the domain. Consequently, even though the outlet of the domain remains open in the simulation, no fluid exits until the time of a mass movement.

We describe these dynamics more mathematically in the following.

First, we can estimate the rate of increase in the volume of the proximal LI as follows:

$$Q_{in} = \frac{dV}{dt} \qquad [S3]$$

where, $Q_{in}$ and $V$ are the inflow and inner volume of the proximal LI, respectively. Given a tube geometry, we can rewrite this equation as follows:

$$\frac{dR}{dt}|_{exp} = \frac{Q_{in}}{2\pi rL} \qquad [S4]$$

where, $R$ and $L$ are the inner radius and length of the proximal LI, respectively. It should be noted here that we assume the diameter of different sections of the proximal LI to be constant, and $\frac{dR}{dt}|_{exp}$ dictates a uniform expansion rate imposed on the entire proximal LI.

At the time of meal intake, and due to the corresponding gastroileal reflexes (see **SI Text 5** below), a massive peristaltic movement occurs, pushing luminal contents towards the distal colon. The duration of this mass movement can be estimated using the balance between the inflow and outflow, $Q_{out}$, of the proximal LI.

$$Q_{in,basal} - Q_{out} = \frac{dV}{dt} \qquad [S5]$$

where, $Q_{in,basal}$ is the basal (off-peak) flow rate at the inlet. It is reasonable to assume that during the short propulsive contractions, $Q_{in,basal}$ is constant. Thus, to solve this equation, we have the following:

$$Q_{in,basal} - (u_{out} \cdot \pi R^2) = 2\pi RL\frac{dR}{dt} \qquad [S6]$$

where, $u_{out}$ is the outflow velocity (which can be assumed to be equal to the velocity of the propulsive contractions: ~0.7 cm/s).[23] In our simulations, we assume a mass movement lasts until it reaches its relaxed volume (corresponding to a relaxed radius of $R_1 = \sim 1\ cm$). Solving Eq. [S4] for the duration of the mass movement, $t_m$, we obtain:

$$t_m = -\frac{L}{u_{out}} \times ln\left(\frac{\pi u_{out}R_2^2 - Q_{in}}{\pi u_{out}R_1^2 - Q_{in}}\right) \qquad [S7]$$

where, $R_2$ and $R_1$ are the radii of the proximal LI at the start and end of the mass movement, respectively. Note that we here neglect the peristaltic nature of the mass movement and assume that the diameter of the proximal LI cannot change as a function of its length. Eq. [S6] can also be rearranged to find the contraction of the proximal LI during mass movement, $\frac{dR}{dt}|_{cont}$, as follows:

$$\frac{dR}{dt}|_{cont} = \frac{Q_{in,basal} - (u_{out} \cdot \pi R^2)}{2\pi RL} \qquad [S8]$$

#### Flow and reaction kinetics

The advection-diffusion equations that are solved for the nutrient, microbes, and FPs can be expressed as:

$$\frac{\partial c}{\partial t} + \nabla \cdot (-D\nabla c + uc) = S \qquad [S9]$$

where, $c$ denotes the concentration/density of either nutrients, FPs, or microbes. $D$ denotes the diffusion coefficient. $u$ is the velocity field that can be calculated from the momentum and continuity equations for an incompressible fluid:

$$\rho \left[\frac{\partial u}{\partial t} + (u \cdot \nabla)u\right] = -\nabla p + \mu \nabla^2 u \qquad [S10]$$

$$\nabla \cdot u = 0 \qquad [S11]$$

where, $\rho$, $\mu$, and $p$ are the fluid density, viscosity, and pressure, respectively. While viscosity strongly varies along the entire LI due to water uptake and with strong dependence on diet composition [4], it is reasonable to assume that most of the 1.5 to 2 L of luminal contents that enter the LI each day have water-like physical properties. We thus take the viscosity and density values of colonic contents to be that of water (values listed in **Table S1**). Regarding pressure, there is a gradual pressure decrease along the flow direction within the simulation domain, but it mostly remains close to ambient pressure, as the colonic contents are assumed to be non-compressible, and the simulation domain changes its volume as a result of inflow or mass movements (outflow).

For the advection-diffusion equations of nutrients, FPs, and microbes (**Eq. [S9]**), the following source terms, $S_n$, $S_{FP}$, and $S_b$ are used, respectively:

$$S_n = -\frac{\lambda}{Y}\frac{c_n}{c_n + K} c_b \qquad [S12]$$

$$S_{FP} = \lambda \varepsilon_{FP} \frac{c_n}{c_n + K} c_b \qquad [S13]$$

$$S_b = \lambda \frac{c_n}{c_n + K} c_b \qquad [S14]$$

Here, $\lambda$, $Y$, and $\varepsilon_{FP}$ are the maximum growth rate, the yield for growth on carbohydrates, and the excretion of FPs, respectively.

The absorption of FPs by the colonic epithelial, $J_{FP}$, is as follows:

$$J_{FP} = J_{FP,max} \frac{c_{FP}}{c_{FP} + K_{FP}} \qquad [S15]$$

with, $J_{FP,max}$, $c_{FP}$, and $K_{FP}$ being the maximum absorption rate, FPs concentration, and absorption constant, respectively.

Since we use a 2D axisymmetric model, the size of the ileocecal valve, $L_{ileo}$, needs to be adjusted such that the area of the revolved ileocecal opening becomes equal to the actual cross-sectional area of the ileocecal opening, $d_{ileo}$. Therefore, an adjusted value is calculated for $L_{ileo}$ and is assumed to be constant throughout the simulations, as follows:

$$L_{ileo} = \frac{d_{ileo}^2}{8R_1} \quad \text{[S16]}$$

#### Boundary conditions, model implementation, and results analysis

A summary of the boundary conditions and initial values used in the simulations are shown in **Fig. S3**. Chosen parameters and model assumptions to describe mixing and flow are further motivated in **SI Texts 4 and 5**. All parameters used in the simulations to specify dynamics and boundary conditions are listed in **Table S1**.

We use COMSOL Multiphysics (Massachusetts, USA) for setting up and execution of the simulation using the software built-in physics. To enhance computational efficiency, we set the maximum time step as 5 s. We used "IF" clauses to implement source terms (e.g., $S_n$) such that, if the concentrations (e.g., $c_n$) fall below a certain threshold (e.g., smaller than $K$), the corresponding source term is assumed to be zero. To ensure that the integrity of our mathematical model is not compromised, we conduct a "test" simulation for the full model scenario (i.e., mass movement, cecum, and varying inflow included). This "test" simulation uses a maximum time-step of 1 s without employing "IF" clauses, and the obtained results are consistent with our other simulations. In all simulations, we select 72 hours (48 hours for FPs simulations), and unless specified otherwise, the figures presented depict results from the final 24 hours to ensure stable dynamics have been reached. Additionally, in choosing between 3D and 2D simulations, we find that a 2D axisymmetric simulation provides the closest representation of physiologically relevant geometry while maintaining computational costs at a reasonably low level. Once the simulations are completed, the data is extracted and then transferred to Python for further analysis and plotting via a custom-written script.

To convert microbial biomass units from optical density to microbial dry weight we assume in this work that 0.5 mg of microbial dry weight is equivalent to $1\ OD600 \cdot mL$.

### 4. Colonic flow and mixing

In gut models, it is often assumed that the flow of food into the LI remains constant throughout the day [16,17]. While such assumptions could yield realistic densities of microbes typically found in feces via implementing further constrains like water uptake , they do not account for the daily fluctuations in microbial densities, especially in the proximal LI, the primary reservoir for gut microbes to persist and grow. Additionally, there is evidence of daily fluctuations in the inflow to the cecum from the small intestine, with peaks occurring after meal intake [4]. This is supported for example by changes in gastrointestinal motor function coordinated with meal intake, which is thought to control the transport of luminal flow through the ileum and ileocecal valve into the LI [24,25]. Accordingly, experimental observations reveal that luminal contents are released by the ileocecal valve in a non-continuous fashion, regulated by the host and depending on digestion activity [24,26]. Consequently, luminal flows are linked to meal timing. To provide realistic inflow conditions, we use measurements of water passage through the ileocecal valve as reported by Phillips and Giller [4] for healthy individuals consuming three major meals a day (**Fig. S4A** with meal intakes shown in red). In our simulations, we specifically assume a daily inflow of 1.5 L through the ileocecal valve constituting a basal flow rate, $Q_{in,basal}$, and peaks in inflow ($10 \times Q_{in,basal}$) starting 1 h after each meal intake (a total of three meals occurring at 7:00, 13:00, and 19:00 hours in each day of simulation) and lasting for 1 h (**Fig. 1C**).

In addition to advection (directed flow), the mixing of colonic contents significantly contributes to the abundance of nutrients, microbes, and growth dynamics. Colonic contents mix as they traverse the LI, resulting from complex interactions between different colonic wall movements. Various methods, including pressure readings and radiological devices, have been used to characterize colonic contractions [27]. These contractions, along with their corresponding pressure waves, can lead to different flow dynamics in various regions of the colon, affecting the spatiotemporal dynamics that influence transit time and local mixing of colonic contents. Depending on the nature, timing, and colonic region where these pressure waves occur, colonic contractions can be either stationary or directional, occurring in antegrade or retrograde directions [28,29].

To account for local mixing within the proximal LI in our simulations, we assume continuous mixing throughout the day by assigning a diffusion coefficient in the advection-diffusion equation representing the mixing condition. To determine a realistic range of mixing parameters (diffusion coefficients) for our model, we first assume that in the absence of wall contractions, the mixing coefficient describes the random movement of microbes in a static fluid, of the order of $\sim 10^{-9}\, m^2/s$, as reported [30,31]. In the presence of colonic wall contractions, the mixing parameter increases significantly. As an upper limit, we assume that under strong mixing conditions, it does not exceed $\sim 10^{-6}\, m^2/s$, as previously estimated [17,18].

The inflow into the cecum from the ileum has been measured at around 1.5 liters per day [4,15]. The majority of this inflow is absorbed by the intestinal epithelium, mainly through water absorption, a major role of the colon. Thus, the net flow rate values could vary along the LI. Since our study primarily focuses on the proximal LI, we disregard this effect and instead assume that the flow rate remains constant throughout the simulation domain but vary over time.

Colonic inflow also brings nutrients, particularly carbohydrates, into the cecum that microbes can use for growth. The types and quantities of nutrients can vary depending on diet, eating behavior, and carbohydrate composition [2,32]. In this study, we assume a total of approximately 0.14

mole of glucose-equivalent carbohydrates enter the cecum after meal intakes each day, a value typical for a British diet consumed in the 1970s (British reference diet) [2].

### 5. Colonic mass movements

Colonic motor activity can be categorized into two types: propulsive and non-propulsive contractions [33]. Non-propulsive contractions primarily involve segmental activity, while intermittent propulsive contractions can move colonic contents over considerable distances, often referred to as mass (or bolus) movements [34]. Both types of contractions can increase following a meal and decrease during sleep [33]. The frequency of mass movements may increase in cases of diarrhea and decrease in constipation, and it can also be influenced by antidiarrheal or laxative drugs [33]. These contractions generate intraluminal pressure waves along the LI, usually propagating peristaltically, followed by bolus movements of colonic contents [23]. In studies involving the constant infusion of external fluids into the healthy human cecum, the movement of fluid out of the ascending colon and its collection from the rectum were found to be intermittent [5,34], underscoring the colon's ability to adapt to the volume of its contents and the occurrence of mass movements, which clear colonic contents in bolus segments (**Fig. S4B**). Although colonic motility exhibits wide fluctuations, these are primarily associated with fasting/eating or sleeping/awakening periods [35]. Colonic pressure waves, constituting mass movement events, are most active in the transverse and left colon and are linked to the urge to defecate [36–38]. However, they can also occur in the proximal colon and partially contribute to colonic flow in that region. In our simulations, we account for colonic pressure waves and presume that flow is primarily governed by mass movements. These mass movements typically occur three to six times a day, often in the morning after waking up and in the late postprandial period, spanning long colonic segments at speeds of approximately 0.5 to 1 cm/s [23,35,37]. Consistent with these findings, we assume in our simulations that after each meal, one mass movement occurs, emptying the proximal LI into the transverse colon at a rate of 0.7 cm/s. This mass movement concludes when the volume of the proximal LI returns to its relaxed state.

### 6. Discussion on additional scenarios

In the main manuscript, we focus on three key physiological parameters and their respective effects on the microbial population: (i) the daily inflow variation, (ii) the presence of a pouch-like cecum, and (iii) the expandable nature of the proximal LI. We refer to the most physiologically relevant model, which includes the cecum, a deformable domain with the expansion-contraction nature of the proximal LI, and varying inflow, as the full model scenario (**Fig. S6**). To further decouple the impact of each of these key parameters, in addition to the different scenarios discussed in **Fig. 1**, we have also simulated additional scenarios (**Fig. S7**). While these scenarios are not physiologically realistic, they offer insights into the role of each parameter.

To characterize the role of the cecum and its influence on growth dynamics, we compare scenarios with and without the cecum in our simulations. When the proximal LI is assumed to be rigid (i.e., non-expandable), none of the combinations, including/excluding cecum and constant/varying inflow, can ensure a stable microbial population (See **Figs. 1B D-E and S8 A-C**). The microbes are washed out in all these cases, with the total microbial mass within the proximal LI approaching zero. However, for an expandable proximal LI, under similar conditions, the likelihood of sustaining microbial populations significantly increases, emphasizing the crucial role of the expandable nature of the proximal LI.

In a scenario with an expandable simulation domain, the presence of cecum can greatly influence microbial densities (See **Fig. S8 D-F**). Specifically, the expandable domain without a cecum, under either constant or varying inflow, can sustain microbes from washout only under high diffusion regimes (See **Fig. S9**). In contrast, with a cecum included this dependency on diffusion becomes less pronounced, and stable microbial populations can be maintained with less mixing. (See **Figs. S6 and S8**). This mainly stems from the dead-end nature of the cecum, where the transport of nutrients/microbes to/from this region relies heavily on mixing. For example, for an intermediate mixing condition (i.e., $D = 10^{-8}\ m^2/s$) and varying inflow with expansion-contraction cycles of the proximal LI volume, we find that the variation in microbial density is governed by two waves of microbial growth: one in the cecum and another in the distal proximal LI (See **Fig. 2D** white arrows).

### 7. Comparison of observed and simulated microbial densities along the LI

Most nutrients that microbes can efficiently digest, including complex carbohydrates, are consumed before leaving the proximal LI. As such, there should be not much more microbial growth beyond the proximal LI. Thus, the daily number of microbes exiting the proximal LI is approximately given by the number of microbes exiting the gut via feces. To estimate the density of microbes at the end of the proximal LI, we can thus, as a rough estimation, take the density of microbes in feces. For instance, microbial densities in the range of 0.001 – 0.1 OD600 and 10 – 100 OD600 have been experimentally measured in the cecum and feces, respectively [39,40]. One crucial factor explaining the difference between the microbial densities found in the cecum and those in feces is the change of concentrations due to water uptake [1,41]. Approximately only 10% of the water/luminal content enters the LI exit via feces, while the rest is taken by the epithelium of the colon, mainly along the distal ascending and the transverse colon. As we do not account for this water uptake in our model, the maximum density of microbes towards the end of the ascending colon in our simulations should be about ten times lower than what is observed in feces. Accordingly, we expect microbial densities to vary between approximately 0.001-10 OD600 in the cecum ascending colon throughout the day. Given the volume of the proximal LI, this corresponds to a variation of the total bacterial mass within approximately > 0.01 mg and 1 g dry weight throughout the day, in line with the simulations of our full model (**Fig. 1F**).

With 1 OD600 being equivalent to approximately $7 \times 10^8$ cells/mL [42], the cell-number densities of microbes within the LI vary within the range of $7 \times 10^5$ to $7 \times 10^{10}$ cells/mL ($7 \times 10^5$ to $7 \times 10^9$ without water uptake), also comparable to ranges obtained in the simulations of our complete model (approximately $10^4$ to $10^{10}$ cells/mL, see **Fig. 4A**). These numbers are also consistent with an upper bound of $\lesssim 4 \times 10^{13}$ cells within the large intestine (or ≈ 25g bacterial dry weight), considering that the ascending colon with lower bacterial densities accounts only for a minor fraction of the total LI volume of approximately 500 mL.
